# Trends in DNA methylation with age replicate across diverse human populations

**DOI:** 10.1101/073171

**Authors:** Shyamalika Gopalan, Oana Carja, Maud Fagny, Etienne Patin, Justin W. Myrick, Lisa McEwen, Sarah M. Mah, Michael S. Kobor, Alain Froment, Marcus W. Feldman, Lluis Quintana-Murci, Brenna M. Henn

**Affiliations:** Department of Ecology & Evolution, Stony Brook University, Stony Brook, NY, United States; Department of Biology, University of Pennsylvania, Philadelphia, PA, United States; Department of Biostatistics, Harvard TH Chan School of Public Health, Boston, MA, United States; Department of Biostatistics and Computational Biology, Dana-Farber Cancer Institute, Boston, MA, United States; Human Evolutionary Genetics, Department of Genomes & Genetics, Institut Pasteur, 75015 Paris, France; Centre National de la Recherche Scientifique, URA 3012, 75015 Paris, France; Center of Bioinformatics, Biostatistics and Integrative Biology, Institut Pasteur, Paris 75015, France; Department of Anthropology, University of California at Los Angeles, Los Angeles, CA, United States; BC Children’s Hospital, Department of Medical Genetics, University of British Columbia, Vancouver, BC, Canada; Institut de Recherche pour le Développement, 75006 Paris, France; Muséum National d’Histoire Naturelle, 75005 Paris, France; Centre National de la Recherche Scientifique, UMR 208, 75005 Paris, France; Department of Biology, Stanford University, Stanford, CA, United States

## Abstract

Aging is associated with widespread changes in genome-wide patterns of DNA methylation. Thousands of CpG sites whose tissue-specific methylation levels are strongly correlated with chronological age have been previously identified. However, the majority of these studies have focused primarily on cosmopolitan populations living in the developed world; it is not known if age-related patterns of DNA methylation at these loci are similar across a broad range of human genetic and ecological diversity. We investigated genome-wide methylation patterns using saliva and whole blood derived DNA from two traditionally hunting and gathering African populations: the Baka of the western Central African rainforest and the ≠Khomani San of the South African Kalahari Desert. We identify hundreds of CpG sites whose methylation levels are significantly associated with age, thousands that are significant in a meta-analysis, and replicate trends previously reported in populations of non-African descent. We confirm that an age-associated site in the gene ELOVL2 shows a remarkably congruent relationship with aging in humans, despite extensive genetic and environmental variation across populations. We also demonstrate that genotype state at methylation quantitative trait loci (meQTLs) can affect methylation trends at some known age-associated CpG sites. Our study explores the relationship between CpG methylation and chronological age in populations of African hunter-gatherers, who rely on different diets across diverse ecologies. While many age-related CpG sites replicate across populations, we show that considering common genetic variation at meQTLs further improves our ability to detect previously identified age associations.

## Introduction

Aging is a degenerative process that is associated with changes in many molecular, cellular and physiological factors. Identifying biomarkers associated with these changes is of great interest to researchers for generating accurate predictions of both chronological and biological age in humans for health care and forensic applications. Recent epigenomic studies have shown that patterns of DNA methylation change substantially with chronological age: genome-wide methylation levels decrease with increasing age, while particular genomic regions, such as CpG islands, become more methylated with increasing age^1-5^. An epigenome-wide study examining over 475,000 CpG sites found significant age-associated changes in DNA methylation at almost one-third of sites^2^, demonstrating the extensive and stereotypic effect of aging on the human epigenome. Many previously proposed molecular biomarkers for aging, including leukocyte telomere length^6^, aspartic acid racemization^7^ and expression levels of certain genes^8-10^, can be challenging to apply for age estimation, due to lack of precision, instability over time, or difficulty in measuring the quantity of interest^11^. In contrast, DNA methylation values measured from relatively few (from three to up to a few hundred) age-associated CpG sites (a-CpGs) have been shown to yield highly precise and accurate estimates of chronological age^12-14^. Recent technological improvements, in particular the introduction of the Illumina Infinium^®^ HumanMethylation450 BeadChip array, have greatly expanded the scope of epigenetics research. This platform increases the density of assayed CpG sites across the human genome compared to the older Infinium^®^ HumanMethylation27 array, leading to the discovery of several novel potential aging biomarkers^15^.

Changes in DNA methylation at putative a-CpGs may be affected both by genetic and environmental factors, in addition to aging itself. Extrinsic environmental factors such as smoking, sun exposure and obesity, for example, are associated with specific changes in DNA methylation patterns^16-19^. Intrinsic factors, such as genetic background, can also influence patterns of epigenetic aging, including ‘baseline’ DNA methylation levels at a-CpGs and the rate of change with age^1, 20, 21^. Importantly, specific genetic variants occurring at different frequencies or involving population specific gene-environment interactions, can lead to patterns of DNA methylation that differ between human ethnic groups^22-24^ and drive divergent patterns of epigenetic aging. Few studies have explored epigenetic aging while also explicitly considering ancestry (but see Zaghlool et al.^25^ and Horvath et al.^26^), and most previous work has focused on cosmopolitan populations of European origin^1, 2, 27^. However, it cannot be assumed that age-related methylation trends identified in one human population will be the same in other populations. Further validation of potential methylation-based aging biomarkers in cohorts of diverse ethnic backgrounds is therefore essential before they can be widely applied in the fields of health care, anthropology and forensics^11^. It is also important to note that different human cell types exhibit significantly different genome-wide methylation patterns^28-31^, a factor that potentially affects a-CpGs as well.

In order to explore the impact of genetic ancestry and cell specificity on epigenetic aging, we methyltyped over 480,000 CpG sites in saliva and peripheral whole blood samples from 189 African hunter-gatherer individuals from two populations: the ≠Khomani San of the South African Kalahari Desert and the Baka rainforest hunter-gatherers (also known as “pygmies”^32^) of the western Central African rainforest. These two populations diverged relatively early from the ancestors of all other modern humans, and exhibit much greater genomic variation than most populations whose global methylation patterns have been assayed so far^33, 34^. The ≠Khomani San, in particular, are among the most genetically diverse populations on Earth^33-35^. Furthermore, the ≠Khomani San and the Baka differ in terms of their nutritional subsistence, ecological environs (semi-desert and equatorial rainforest, respectively) and physical activity levels from the widely studied cohorts of cosmopolitan populations. By using methyltype data from these populations, we are able to explore patterns of epigenetic aging across a greater range of human genetic diversity and test previously published methods of estimating epigenetic age in order to determine their accuracy across ethnicities and cell types.

## Materials and Methods

### DNA and ethnographic collection

Saliva was collected from 56 ≠Khomani San individuals (aged 27-91, median age 62) and 36 Baka individuals (aged 5-59, median age 30) using Oragene DNA self-collection kits (Figure S1). Blood was collected from 97 additional Baka individuals (aged 16-90, median age 44) for a previous study^24^ (Figure S1). DNA samples from the ≠Khomani San were collected with written informed consent and approval of the Human Research Ethics Committee of Stellenbosch University (N11/07/210), South Africa, and Stanford University (Protocol 13829), USA. ≠Khomani San participant ages were verified ethnographically on a case-by-case basis. Various documents, such as birth certificates, wedding certificates, school records, and other forms of identification (e.g. apartheid government IDs), were cross-referenced to identify any inconsistencies. Local major events, such as the creation of the Kalahari National Park in 1931, were also used to verify participant’s life history stage. DNA samples from the Baka were collected with informed consent from all participants and from both parents of any participants under the age of 18. Ethical approval for this study was obtained from the institutional review boards of Institut Pasteur, Paris, France (RBM 2008-06 and 2011-54/IRB/3). Baka participant ages were determined ethnographically by Alain Froment by comparing individuals from a single cohort to one another and with reference to major historical events. Baka individual ages are estimated to be accurate to within five years. The Baka saliva sample was known to contain nine trios and nine unrelated individuals.

### DNA methylation data generation and data processing

The 97 Baka whole blood samples were previously processed and published in Fagny et al.^24^while the 89 saliva samples were newly generated for this study. DNA extracted from all samples was bisulfite converted, whole genome amplified, fragmented and hybridized to the Illumina Infinium^®^ HumanMethylation450 BeadChip. This array assays methylation levels at over 485,000 CpG sites throughout the genome through allele-specific single-base extension of the target probe with a fluorescent label. The saliva samples from both populations were methyltyped together in two batches, and the blood samples in six batches. Methylation data from Illumina methylation arrays often exhibits substantial batch effects; that is, samples from one run may vary systematically from the same samples on a different run due to technical artefacts. In order to account for this, we included one ≠Khomani San individual from the saliva dataset on both runs. The overall correlation between beta values from the first and second runs was over 0.99, indicating that batch effects are relatively minimal in our saliva dataset. Technical replicates were also included in the whole blood methyltyping, and the overall correlations of beta values between repeat individuals were all greater than 0.98. The intensity of fluorescence was used to calculate methylation levels. Probes with a detection p-value above 0.01, those that were found to map to multiple genomic regions or to the sex chromosomes, or to contain known SNPs were removed, leaving 334,079 sites in the saliva dataset and 364,753 sites in the blood dataset for subsequent analysis. Probe SNPs were identified using the 450k array annotation file published by Price et al.^36^ and by cross-referencing the genomic coordinates of our samples’ genotype data and the methyltyping probes using bedtools. These values were background and colour-corrected, and technical differences between Type I and Type II probes were corrected by performing quantile and subset-quantile within-array normalization (SWAN) using the *lumi* and *minfi* R packages. For a discussion of the various technical issues inherent in the 450k array design, see Dedeurwaerder, S. et al.^37^ and Makismovic, J. et al.^38^. One Baka individual had abnormally low bisulfite controls, which made methylation values for that sample unreliable. We conducted principal component analysis (PCA) separately on the saliva and blood methylation datasets using the prcomp function in R. A biplot of the first two principal components revealed that the previously flagged Baka individual and six ≠Khomani San individuals were extreme outliers (Figure S2), and these samples were excluded from further analyses. All analyses were performed using continuous beta values for each CpG site, which range from 0 (indicating that the site is completely unmethylated) to 1 (completely methylated).

### Single nucleotide polymorphism (SNP) genotype data

The DNA samples were genotyped on either the Illumina OmniExpress, OmniOne or 550k SNP array^24, 35, 39, 40^. All Baka individuals and 48 of the ≠Khomani San individuals were successfully genotyped. OmniExpress data from the Baka blood samples was imputed using the results of the OmniOne genotyping. The datasets were filtered using a genotyping threshold of 0.95 and a minor allele frequency threshold of 0.01.

### Ancestry inference

We intersected the genotype data generated for the Baka and ≠Khomani San with genotype data from African and European populations (specifically the Biaka pygmies, Mbuti pygmies, Namibian San, southern Bantu-speakers, Kenyan Bantu, Yoruba, French and Italian) generated by the Human Genome Diversity Project (HGDP) on the Illumina HumanHap array^41^. We performed an unsupervised clustering analysis using ADMIXTURE^42^ on the resulting dataset of 254,080 SNPs from 319 individuals in order to determine the global ancestry proportions. We specifically estimated the genetic contributions from Bantu-speaking agriculturalists and Europeans to the hunter-gatherer populations. Prior work has demonstrated an average 6.5% of ancestry from neighbouring Bantu-speakers in the Baka population^40^, and an average of 11% for each of Bantu and European ancestry in the ≠Khomani San^35^.

### Saliva epigenome-wide association study (EWAS)

We used the R package CpGassoc to conduct an epigenome-wide association test (EWAS) for the Baka saliva data. Family identity as a fixed effect, the first two methylation PCs, and percentage of Bantu ancestry were used as covariates in testing for association with age. We used the program EMMAX with the dosage option to conduct the EWAS on the ≠Khomani San methylation data. After removing the outlier individuals, we generated a Balding-Nichols kinship matrix using genotype data from the remaining 44 individuals, which was included in the model to correct for relatedness within the population. For the ≠Khomani San analysis, proportions of European and Bantu ancestry, methyltyping batch and the first PC were used as covariates. The combination of ancestry and PC covariates used in the EWAS was selected in order to minimize the genomic inflation factor (Figure S3). We note that these low genomic inflation factors were obtained by including PCs that were moderately correlated with age as covariates (Figure S4), which may decrease our power to detect a-CpGs. By minimizing the genomic inflation factor in this way, our EWAS are likely to be overly conservative, especially given that a substantial fraction of the 450k array becomes differentially methylated with age^1, 2, 43^. CpGassoc was used on the Baka saliva dataset because it allowed family identity to be included as a fixed covariate, which produced the lowest overall λ. This may be due to the fact that the kinship matrix does not account as effectively for the presence of many first-degree relatives in this dataset. We applied a Benjamini-Hochberg corrected threshold to both EWAS to identify CpG sites whose methylation levels vary significantly with age at a FDR of 5%.

### Blood EWAS

We performed a correction for cell-type composition using the method described by Houseman et al. implemented in the *minfi* package^44^. This compares the observed methylation data from the Baka with reference profiles of each cell type. Proportions of these cell types can vary significantly with age, and, because each cell type has a distinct methylation profile, it is important to correct for heterogeneity in order to avoid spurious correlations between methylation and age in whole blood^45^. We used EMMAX with the dosage option to conduct the analysis on the Baka whole blood methylation data. Genotype data was used to generate a Nichols-Balding kinship matrix of all the individuals, which was included in the model to correct for unknown relatedness within the population. The proportion of Bantu ancestry, methyltyping batch, first three PCs, as well as the estimated proportions of five blood cell types (CD8 T lymphocytes, CD4 T lymphocytes, B lymphocytes, natural killer lymphocytes and monocytes) were used as covariates. The combination of ancestry and PC covariates used in the EWAS were selected in order to minimize the genomic inflation factor (Figure S3). We applied a Benjamini-Hochberg corrected threshold to the Baka blood EWAS to identify CpG sites whose methylation levels vary significantly with age at a FDR of 5%.

### Meta-analysis

We conducted a meta-analysis by combining p-values from both ≠Khomani San and Baka saliva EWAS using Fisher’s method^46^. We applied a Benjamini-Hochberg corrected threshold to the Fisher’s p-values to identify CpG sites whose methylation levels vary significantly with age at a FDR of 5%.

### Hyper- and hypomethylation with age

For every significant CpG site identified in both the saliva EWAS, we fit a linear model of methylation level with age using the R function lm and calculated the slope to determine if it exhibited a hypermethylation (positive slope) or hypomethylation (negative slope) trend with age. We also calculated the residual standard error, multiple r^2^ and AIC value (using the R function AIC) of the linear model. We then fit a new model after first log transforming the age, and recalculated the residual standard error, multiple r^2^ and AIC value. For every site, we then calculated the difference in the residual standard error, multiple r^2^ and AIC value between the linear model and the log-linear model. For all three of these measures, we performed a t-test to compare sites that become hypermethylated with age to those that become hypomethylated with age. Note that, under AIC, models are only considered a significantly better fit if the difference in AIC values is greater than 2^47^. However, as our goal was to examine general trends in the characteristics of hypermethylated and hypomethylated sites, we included the entire distribution of AIC value differences in our analysis.

### Replication of previous studies

We compiled a comprehensive list of 163,170 significant a-CpGs published from 17 studies of methylation and aging conducted in any tissue type^1-3, 13-15, 25, 27, 48-56^. We compared the significant a-CpGs we identified in our three EWAS and the meta-analysis to this list and found 107 a-CpGs that were uniquely identified in our study. For the dataset in which a given novel a-CpG site was identified, we fit a linear model of methylation level and age using the R function lm to determine the slope of the relationship and therefore the direction of the trend with age (hyper- or hypomethylated). We also calculated the Pearson’s correlation coefficient between methylation and age using the R function corr.

### Age prediction

We applied a previously published multi-tissue epigenetic age calculator to estimate the chronological ages of our sampled individuals^12^. The calculator accepts methylation array data as input and outputs a DNA methylation-based age estimate. We used datasets that were not filtered for any probes because the normalization step of the algorithm would not run with a large quantity of missing data. We also tested 450k methylation data from a total of 60 European individuals that were freely available from the Gene Expression Omnibus data repository (GSE30870^3^ and GSE49065^43^). We plotted the estimated age against the individual’s self-reported age and compared the line of best fit to the data with line x=y, which represents perfect prediction.

### Methylation quantitative trait loci (meQTL) scan

We identified *cis*-meQTLs in the Baka blood samples by conducting linear regressions in R of the methylation value at each of the 346, 753 CpG sites against the genotype dosage of all SNPs that lay within 200kb of that site, and had a minor allele frequency of at least 10% in the sample. 11,559 significant *cis*-meQTL associations were identified by applying a Benjamini-Hochberg correction to the p-values at a FDR of 1%, as determined by 100 permutations.

### Conditional analysis

We performed a conditional association analysis for each a-CpG with a significant meQTL by including the genotype state of the associated SNP as an additional covariate in the model. Because EMMAX cannot handle missing values, and because some genotype information was missing, we repeated the ‘baseline’ EWAS for the Baka whole blood data and performed the conditional analysis using CpGassoc correcting for all the same covariates, but excluding the Balding-Nichols kinship matrix. We also performed a permutation analysis by pairing each CpG site with a randomly chosen meQTL SNP and repeating the conditional EWAS, where the genotype state of the ‘false meQTL’ was included as a covariate instead of the ‘true’ one. We permuted CpG-SNP associations a total of 100 times to build a distribution of effects of a random meQTL on general age-association trends.

## Results

### Principal component analysis and ADMIXTURE

We performed principal component analyses (PCA) to determine if there were factors other than age driving systematic differences in DNA methylation profiles. PCA were conducted on the saliva and blood datasets separately, since it is expected that these tissues will differ substantially in their methylation profiles^29^. Individuals clustered together by batch identity in biplots of the first and second PCs, demonstrating that batch effects were the strongest drivers of DNA methylation profile differences, as expected^57^, but population identity (for the saliva dataset) and sex did not appear to drive clustering in the first two PCs (Figure S2). Six ≠Khomani San and one Baka individual were removed from subsequent analyses because their methylation profiles were extreme outliers. We found a significant correlation for some PCs with age, in particular saliva PC 1 with ≠Khomani San age and blood PCs 1 and 2 with Baka age (Figure S4).

Both the Baka and ≠Khomani San have experienced recent gene flow, to differing extents, from Bantu-speaking agriculturalists and additionally, for the ≠Khomani San, with Europeans^40, 58-60^. Since DNA methylation patterns vary substantially across human populations, it is possible that ancestral makeup could also affect patterns of epigenetic aging in admixed individuals^22, 23^. To account for this, we inferred global ancestry proportions using ADMIXTURE for a total of 181 individuals for whom we had SNP genotype array data. Based on previous studies of African genetics, we expect distinct ancestral components corresponding to Pygmy, San, European and Bantu-speaking populations to be present to some degree in our dataset^35, 39, 40^. Therefore, we assumed *k*=4 ancestries when running the ADMIXTURE algorithm (Figure S5). Both the ≠Khomani San and the Baka populations remain relatively endogamous, and coupled with field sampling bias, members of extended families are often collected together. Therefore, we also used the genotype data to generate Balding-Nichols^61^ kinship matrices for the association analyses of the ≠Khomani San saliva and the Baka blood datasets to control for the degree of relatedness between individuals in subsequent analyses. Genetic relationship matrices have been shown to appropriately control for stratification in association studies^62^.

### Epigenome-wide association studies

We conducted an epigenome-wide association study (EWAS) of DNA methylation level and chronological age in each of the three datasets: the ≠Khomani San saliva, the Baka saliva, and the Baka blood. We identified 399 CpG sites in the Baka saliva, 276 sites in the ≠Khomani San saliva and 306 sites in the Baka blood that were significantly associated with age at a false discovery rate (FDR) of 5% (Figure 1, Tables S2-4). 67 of these sites replicated independently in both saliva EWAS and 26 in all three EWAS (Figure 2).

**Figure 1.**
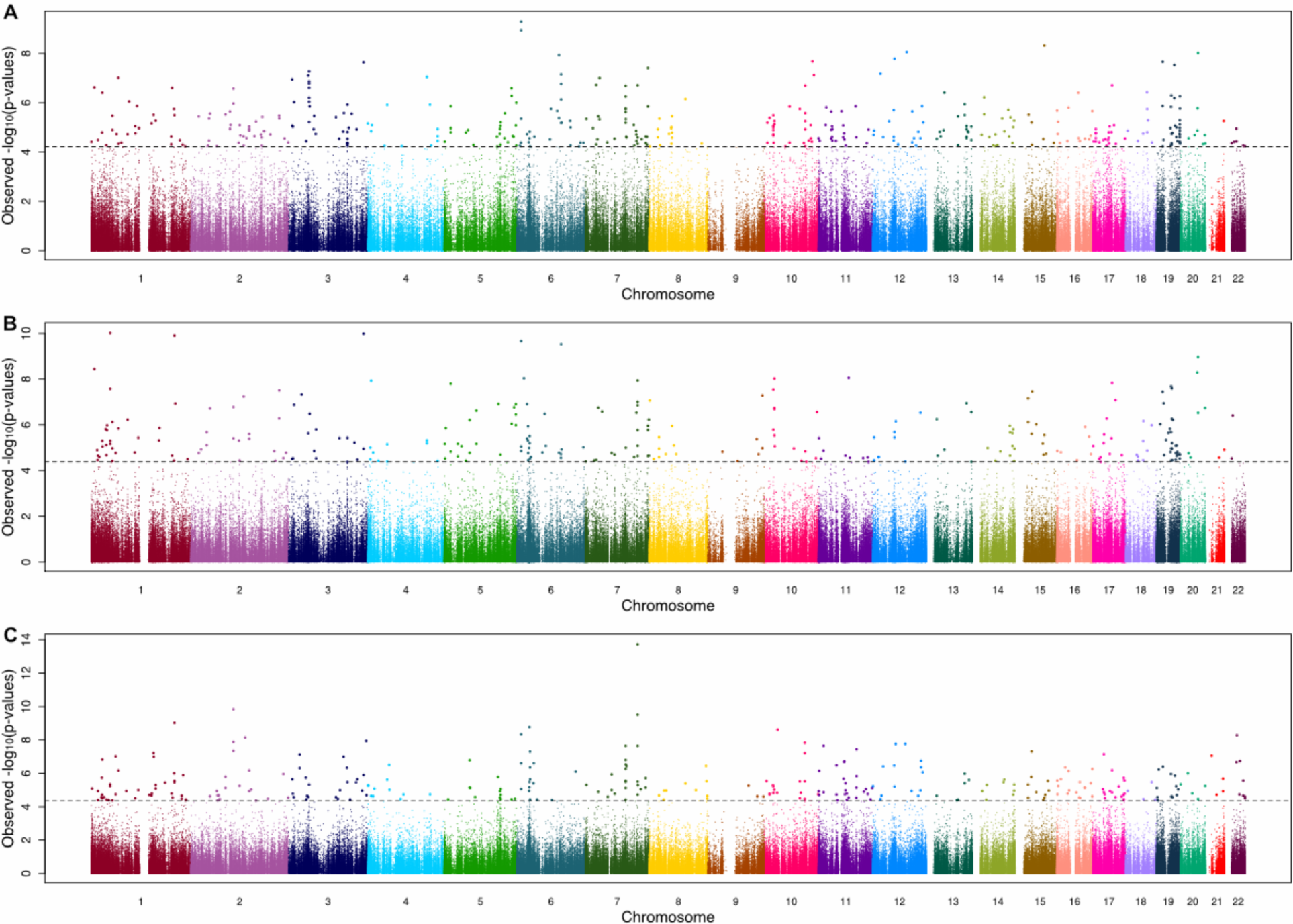
Manhattan plot of epigenome wide association study (EWAS) for age associated CpGs. The —log_10_ p-values from the EWAS are plotted against the assayed autosomal genomic CpGs for A) the Baka saliva dataset, B) the ≠Khomani San saliva dataset and C) the Baka blood dataset. All samples were assayed on the Illumina InfiniumR HumanMethylation450 BeadChips. The horizontal dashed line in each panel represents the Benjamini-Hochberg-corrected threshold for significance (FDR of 5%) for each EWAS.

**Figure 2.**
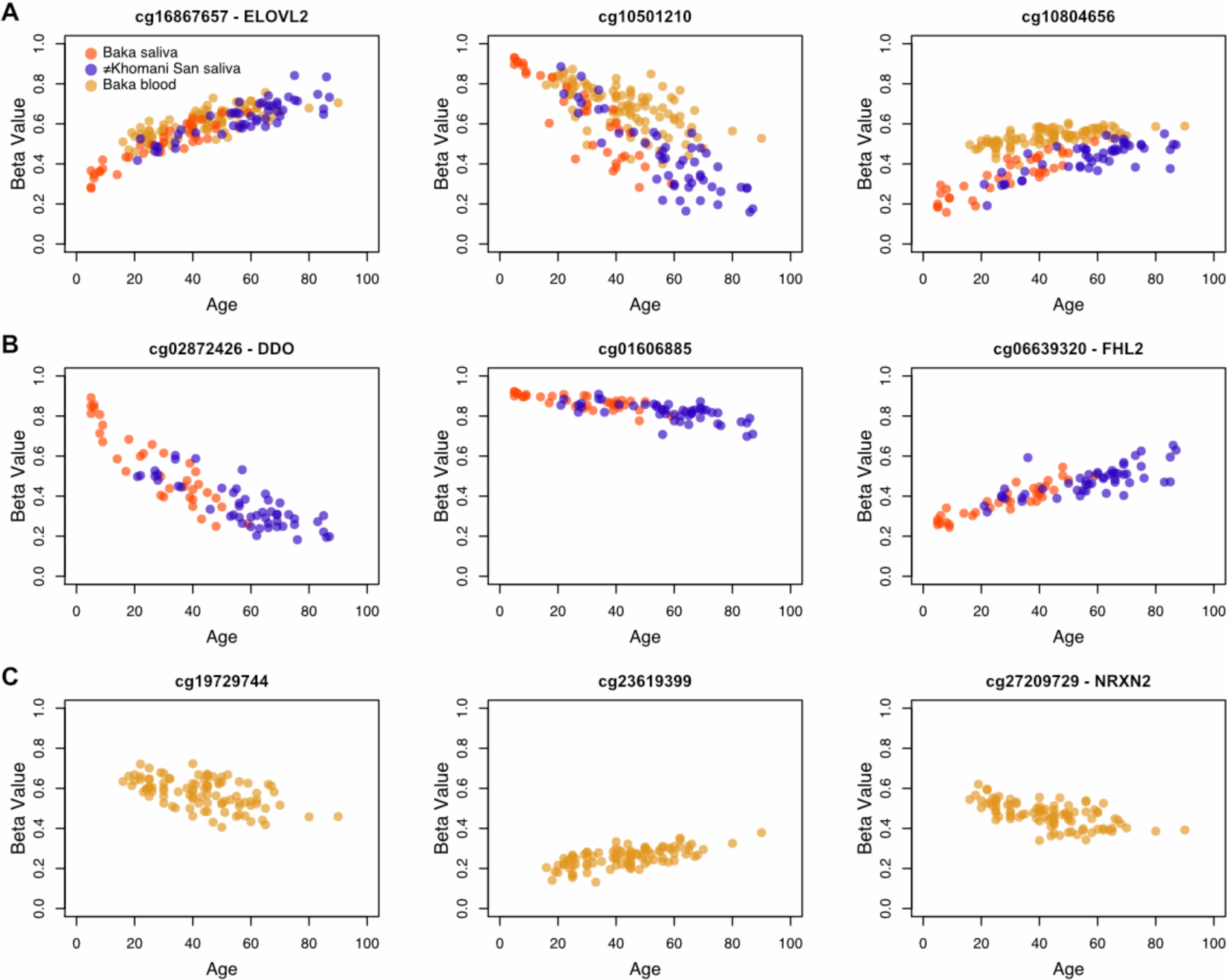
Scatterplots of beta value versus age for a-CpGs. Methylation levels as beta values, which are continuous from 0 (indicating that the site is completely unmethylated) and 1 (indicating that the site is completely methylated), are plotted against age for three of the age-associated CpG sites (a-CpGs) that were identified as significant in A) all three epigenome-wide association studies (EWAS), B) only the two saliva EWAS, and C) only the Baka blood EWAS. Beta values plotted here are not adjusted for the covariates included in each EWAS.

### Meta-analysis of saliva EWAS

In order to improve our power to detect significant age associations in hunter-gatherer saliva, we performed a meta-analysis by calculating Fisher’s p-values from the p-values of both saliva EWAS. We identified 2060 CpG sites that were significantly associated with age at a FDR of 5% in the meta-analysis of our saliva studies. Of these, 1500 (72.8%) show a hypermethylation trend (increasing beta value) with age and 560 (27.2%) show a hypomethylation trend (decreasing beta value) with age (Table S5). The location of each of these a-CpG sites relative to specific genes, genic features and CpG islands was determined from the 450k annotation file available from Illumina. Among these a-CpGs, 1250 (60.7%) fall in CpG islands, 80 in island shelves (3.9%; 50 ‘North’ and 30 ‘South’), 413 in island shores (20.0%; 239 ‘North’ and 174 ‘South’) and 317 (15.4%) in ‘open sea’. When considering all CpG sites assayed in the saliva EWAS, 33.2% of them fall in CpG islands, 9.0% in shelves, 23.6% in shores and 34.2% in open sea. All but 36 of the island sites (2.9%) showed a hypermethylation trend with age, while 73 ‘open sea’ sites (23.0%) showed a hypomethylation trend with age, which is broadly in line with previously reported trends (Figure 3A)^2, 3, 31^. 1536 of our a-CpGs were annotated to specific genes. Among these, we counted the number of sites in each of the following six genic regions: 1^st^ exon, 3’ untranslated region (3’ UTR), 5’ untranslated region (5’ UTR), gene body, within 1,500 base pairs of the transcriptional start site (TSS) and within 200 base pairs of the TSS (Figure 3B).

**Figure 3.**
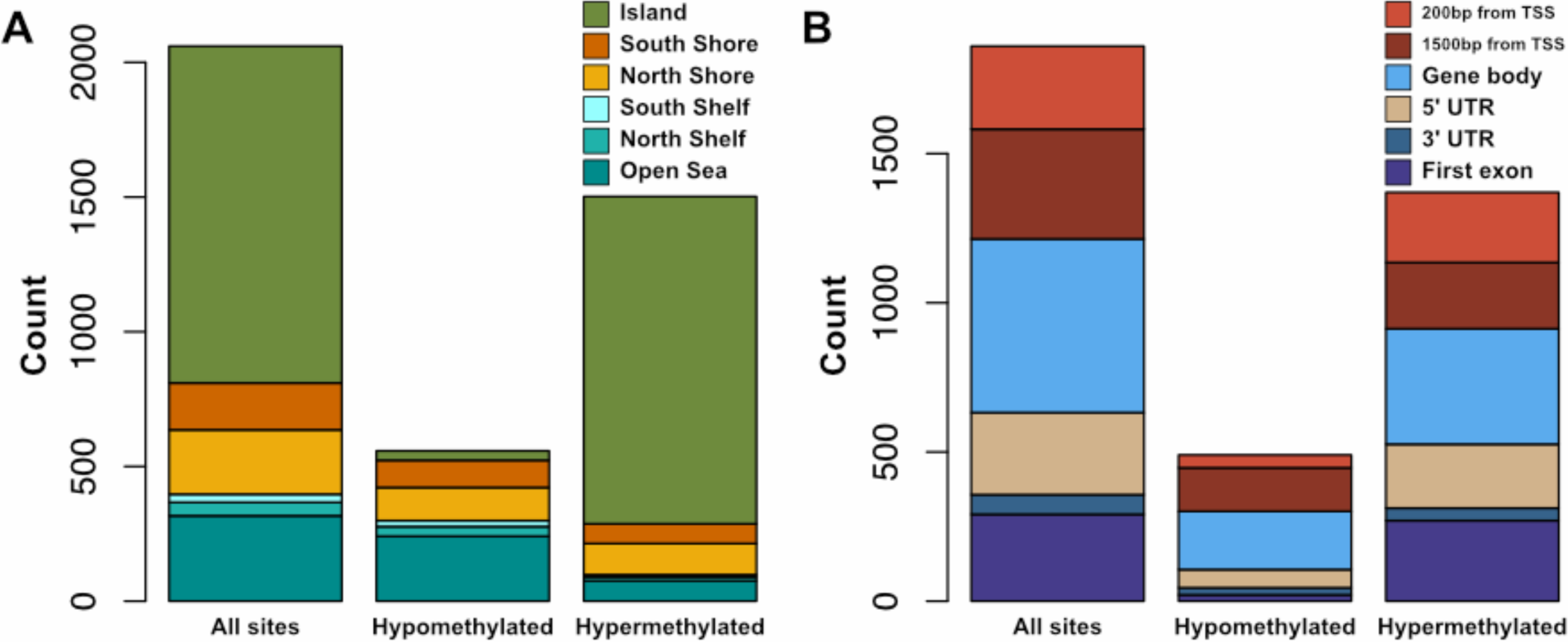
Locations of a-CpGs according to genic features and hyper-/hypomethylation trend. Bar plots indicate counts of a-CpGs and their physical positions relative to A) CpG islands B) genes. In cases where a single CpG site was annotated to multiple gene regions, each region was counted separately. Annotations were provided in the probe information file from Illumina. 76% of a-CpGs become hypermethylated (increase in beta value) with age, and the majority of these lie in CpG islands. By contrast, none of the a-CpGs that become hypomethylated (decrease in beta value) with age lie in CpG islands

We noted that several a-CpGs exhibited a log-linear change in methylation level with age, and particularly in children, as previously reported by Alisch et al.^48^. Interestingly, we observed this pattern more frequently in a-CpGs that become *hypomethylated* with age. We systematically tested this observation by fitting a linear model to the beta values at each of these 2060 sites for both direct chronological age and a log-transformation thereof, and calculated the residual standard error and the multiple r^2^ values for both models. We found that these two classes of sites showed significantly different distributions of residual standard error (p = 1.38 × 10^−36^) and multiple r^2^ (p = 6.14 × 10^−86^). We also calculated the difference in Akaike Information Criterion (AIC) values between the linear and log-linear models of methylation level and age^47^. We then performed a t-test on the difference in AIC values for linear and log-linear models and found, again, that sites that become hypermethylated with age are significantly different from sites that become hypomethylated (p = 1.95 × 10^−76^). All three methods yielded the same general trend: hypomethylated sites tended to be better fit by log-linear models, as demonstrated by their generally higher r^2^ values, lower residual standard errors and lower AIC values when fit by a log-linear rather than a strictly linear model, while hypermethylated sites tended to be better fit by linear models (Figure 4).

**Figure 4.**
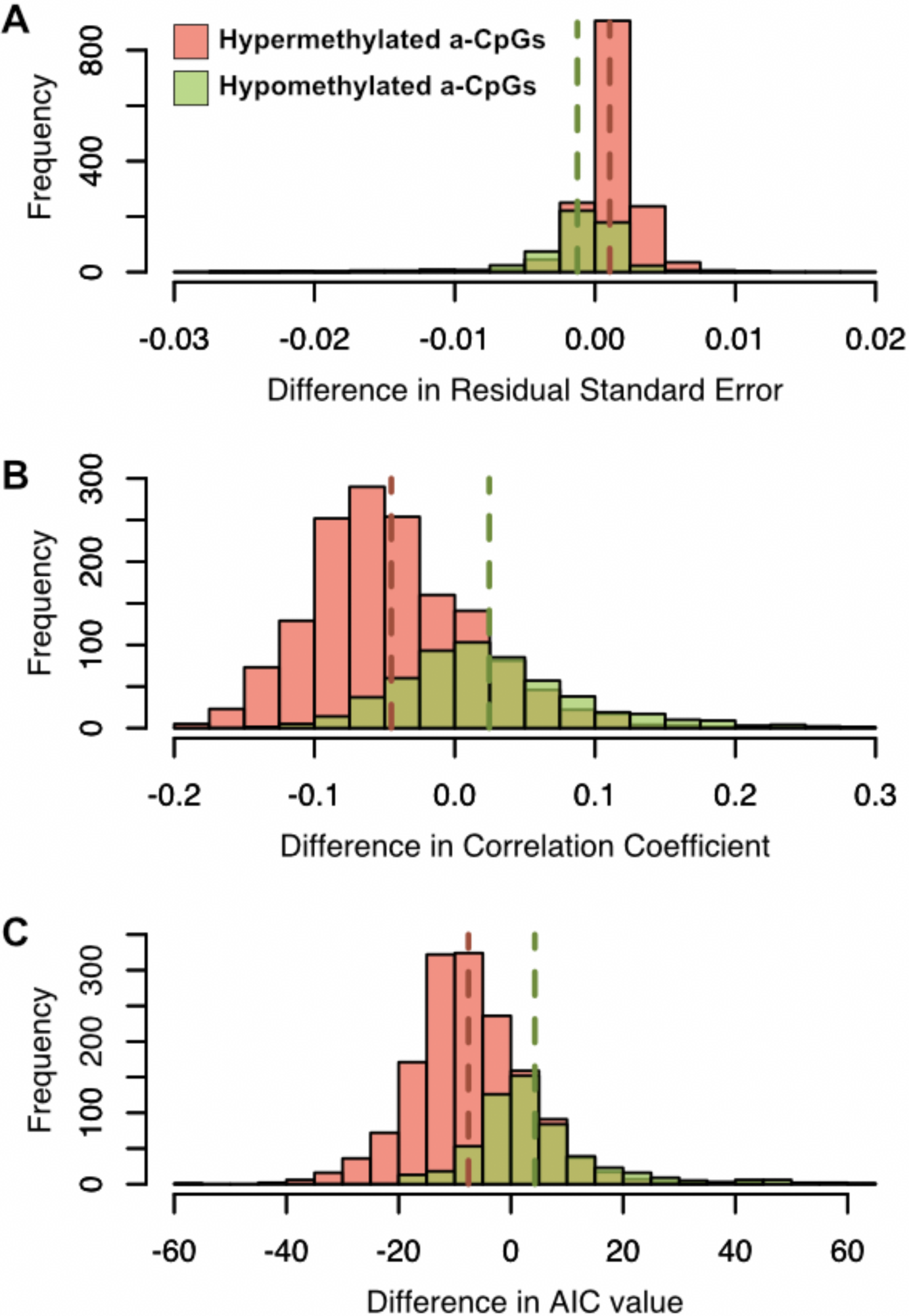
Evaluating the fit of log-linear vs. linear models of methylation level and age to hyper- and hypomethylated a-CpGs. Each of the 67 significant a-CpGs that were identified in both saliva EWAS were fit with both a linear and log-linear model of age with methylation level. The distribution of the differences in A) residual standard error B) correlation coefficient and C) AIC values between the linear model and the log-linear model are shown. The means of the distributions are indicated by dashed vertical lines of the same colour. The linear model is a better fit for the relationship between methylation and age when differences in residual standard error are large and positive and when differences in correlation coefficient and AIC value are large and negative. By all three measures, a-CpGs that hypermethylate with age (orange) are better fit by a linear model and a-CpGs that hypomethylate (green) by a log-linear model.

### Replication of previous studies

We sought to determine the independent replication rate of significant a-CpGs that we identified by searching the literature for studies that quantitatively investigate the relationship between CpG methylation and chronological age. We included 17 studies conducted on either 27k or 450k array technologies in any human tissue^1-3, 13-15, 25, 27, 48-56^. We found that over 95% of the a-CpGs sites we identified in our analyses were reported in one of these previous studies. However, we also found 107 a-CpG sites that were uniquely identified in our study of African hunter-gatherer groups. For each of these 107 sites, we calculated the Pearson’s correlation coefficient between beta value and chronological age and also fit a linear model to determine the slope and trend of the association (Table S1). In order to identify robust aging markers from among these novel a-CpGs, we focused on sites that either exhibited a high Pearson’s correlation value (absolute value over 0.6) or a high slope (absolute value over 0.001 beta value per year); 19 of these 107 a-CpGs met at least one of these two criteria (Figure 5).

**Figure 5.**
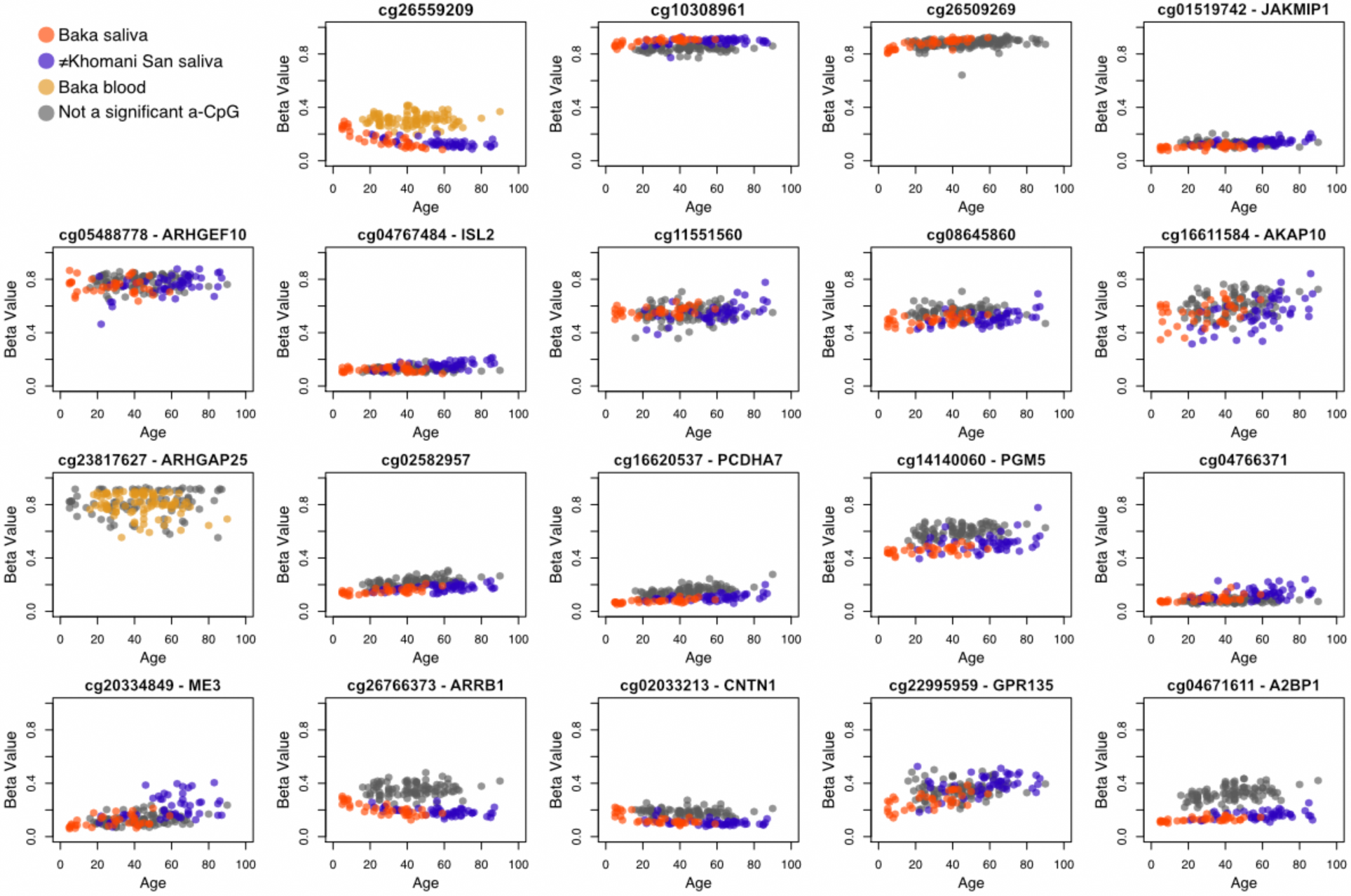
Scatterplots of methylation level and age for novel age-associated CpG sites. Methylation (beta) value is plotted against age for the 19 novel a-CpGs exhibit either an absolute Pearson correlation coefficient of 0.6 or higher, or an absolute slope of 0.001 or higher. Novel a-CpGs were identified if they exceeded the Benjamini-Hochberg-corrected threshold for significance (FDR of 5%) in at least one of the three EWAS or the metaanalysis conducted in this study. Beta values plotted here are not adjusted for the covariates included in each EWAS. Points are greyed out if the CpG site was not identified as significantly associated with age in that dataset.

The site cg16867657, annotated to the gene *ELOVL2,* is significantly associated with chronological age in all three datasets, across populations and tissues. This site was first identified as a potential biomarker for age by Garagnani et al.^15^ and replicated in subsequent epigenetic aging studies of additional cohorts of European, Hispanic and Arab descent^1, 2, 25, 27^. By observing a signal of age association independently in three African cohorts following different lifestyles, and using DNA sourced from two different tissue types, we further validate the use of cg16867657 methylation as a true biological marker for age across the full spectrum of human diversity. The pattern of age-related methylation change is also remarkably congruent across blood and saliva^15, 63^.

In the saliva datasets, we observed a significant age-associated hypomethylation signal in the transcriptional start site of the gene D-aspartate oxidase *(DDO)* at cg02872426 in both African populations (Figure 2B). This site was previously identified in a study of whole blood of Arab individuals^25^ and other CpG sites annotated to *DDO* have also been previously associated with age^25, 53, 64^. We identified additional sites (cg00804078, cg06413398 and cg07164639) in the transcriptional start site of the gene *DDO,* which exhibit hypomethylation with age at a relaxed significance threshold of p < 0.001 in all three datasets (Figure S6).

### Testing an epigenetic aging predictor

DNA methylation can be affected by genetic variation, as well as environmental and lifestyle variation during development. We therefore asked how accurately existing age prediction models, developed primarily on methylation data derived from individuals of European ancestry, would perform on our African datasets. We applied a multi-tissue age predictor developed by Horvath^12^ to all three datasets, hereafter referred to as the “Horvath model” (Figure 6A). This model uses a linear combination of methylation information from 353 sites, termed ‘clock-CpGs’, to produce an estimate of age. The DNA methylation age estimates for the Baka saliva were very accurate, with a median absolute difference of 3.90 years between the true and estimated ages (r = 0.94), and the estimates for the ≠Khomani San saliva dataset had an overall greater median absolute difference of 6.01 years (r = 0.90) (Figure 6B), typically underestimating the chronological age. In order to investigate whether the reduction in accuracy was specific to the ≠Khomani San, we applied the age predictor to European methylation datasets from blood (Gene Expression Omnibus datasets GSE30870^3^ and GSE49064^43^). We observed a similar underestimation of age in older Europeans, suggesting that underestimation in adults older than 50 years is not indicative of a ≠Khomani San-specific slowdown in the epigenetic aging rate (Figure 6C).

**Figure 6.**
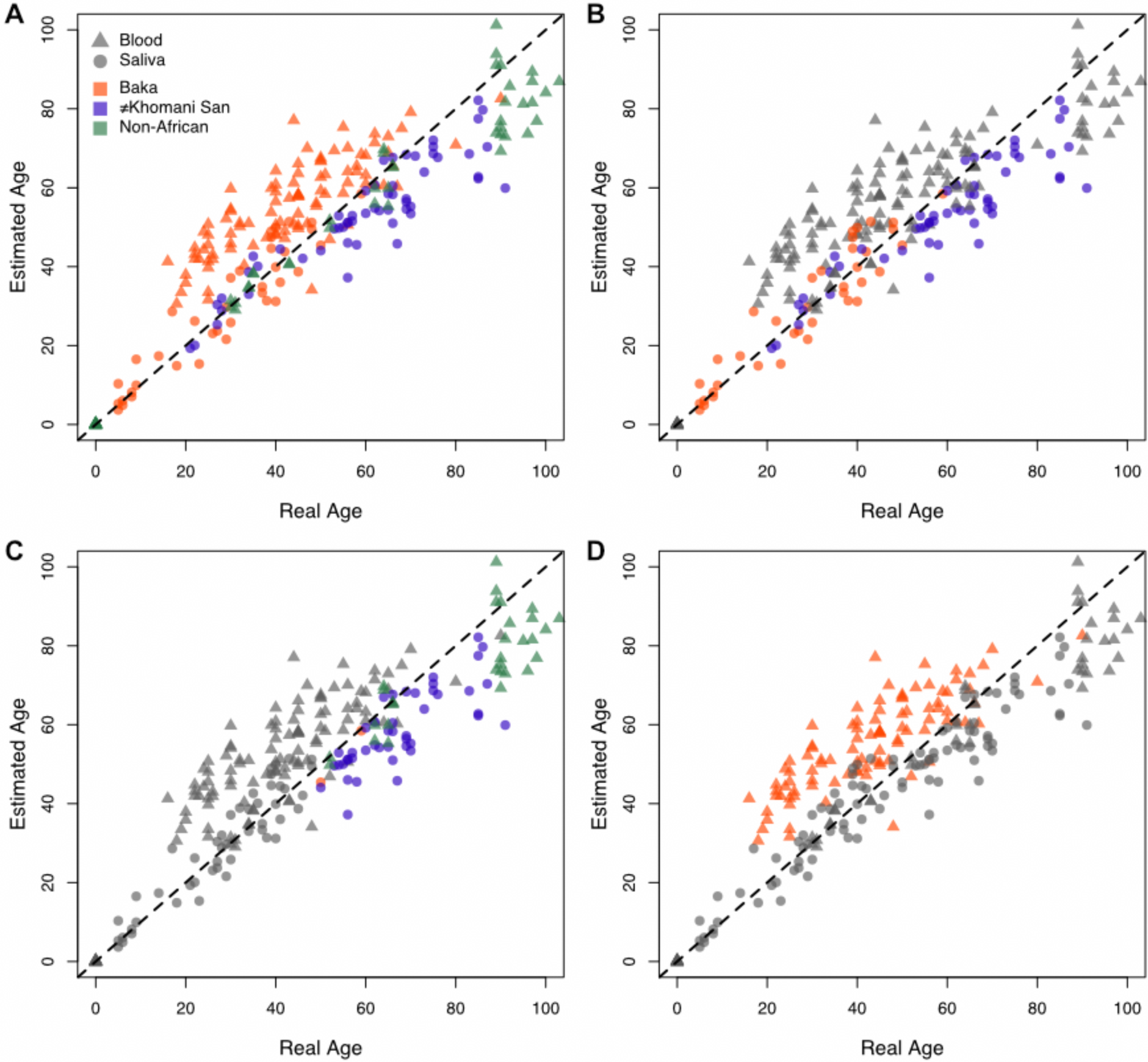
Scatterplots of true age against estimated age as predicted by the Horvath model. Chronological age reported by individuals is plotted against estimated ages generated from epigenetic data using Horvath’s age prediction model^12^. All four panels show the age estimates for the Baka, ≠Khomani San and European blood and saliva datasets, coloured to emphasize different features of the data. Blood and saliva tissue sources are indicated by triangles and circles, respectively. A) All data points, coloured based on population identity. B) Only estimates for African saliva data are coloured; all others are greyed out. C) Only estimates for individuals whose true age is 50 or more, except for the Baka blood data, are coloured. D) Only estimates for Baka blood are coloured. The dashed line represents perfectly accurate prediction of chronological age.

Finally, we observed a systematic overestimation of age from the DNA methylation profiles of Baka blood (median absolute difference of 13.06 years, r = 0.81) (Figure 6D). It is important to note that the correlation between chronological and estimated age remains high, and the discrepancy between the two may be indicative of technical artefacts or batch effects in the application of the arrays. However, it is not possible to rule out a biological driver that causes Baka blood to exhibit increased epigenetic age under the Horvath model (see Discussion).

### Methylation quantitative trait loci in age-related CpG sites

Given the observed differences in accuracy of age estimation in different human populations, we sought to further understand why age-related epigenetic patterns might not replicate across study cohorts. Even at lower significance thresholds, success in reconciling reported epigenetic signals of aging in different studies has been mixed^1^. Indeed, several age-related CpG sites that have been reported previously did not replicate in our populations. There are many potential reasons for this, including our smaller sample sizes, and the comparison of different tissue types which may exhibit tissue-specific patterns of methylation with age. However, it is also possible that population-specific genetic variants or different allele frequencies may drive these differences.

In order to explore this, we sought to determine if methylation quantitative trait loci (meQTLs) play a role in these disparate aging patterns. MeQTLs are genetic variants that are statistically associated with methylation levels at distant CpG sites^65^. We scanned all 346, 753 CpG sites in the Baka blood dataset for *cis-meQTL* associations, where a SNP was considered in *cis* if it was within 200kb of the CpG site. We fit a linear model of methylation by genotype state using chronological age, sex and blood cell type proportions as covariates. We identified 11,559 meQTLs at a FDR of 1% in the Baka blood dataset. We also compiled a list of 18,229 a-CpGs identified in previous studies of blood methylation^1, 3, 14, 15, 25, 27, 48-52, 54-56^. Interestingly, there is an overlap of 901 CpG sites that were identified as being associated with age in Europeans and are also associated with a specific *cis* genetic variant in the Baka. This is more overlap than would be expected by chance, as determined by randomly sampling and intersecting 18,229 CpG sites with the 11,559 significant meQTLs; after 10,000 simulations, the maximum overlap obtained under this null scenario was 662 (p < 0.001). Only eight of these 901 sites were among the 306 significant a-CpGs identified in our EWAS of Baka blood methylation.

We performed a conditional analysis to determine if incorporating genotype information recovers significant age association in the Baka at these CpG sites. For each of these 901 CpG sites, we included the genotype state at the associated meQTL as an additional covariate and repeated the association analysis. We also permuted all the CpG-SNP associations 100 times by assigning each of the 901 CpG sites a SNP selected at random from among the 11,559 identified meQTLs. In the true conditional analysis, we observed an overall upward shift in the distribution of −log 10 p-values when meQTL-specific genotype data was included (Figure 7A), indicating that incorporation of the meQTL genotype generally improves the age-methylation association for these CpG sites (mean increase in −log 10 *p*-value after conditional analysis of 0.15); this increase was not observed in our permutation analysis when a random SNP covariate was included (mean difference in −log 10 p-value of -0.008) (Figure 7B). More specifically, 39 of the 901 CpGs (4.3%) that were not significantly associated with age at a FDR of 5% in the original EWAS recovered significance when true meQTL genotype was included, while this occurred only 0.17% of the time in the permutation analysis. We observe that a small number (15 out of 901, 1.7%) of CpGs decrease in significance by over one order of magnitude, which may be due to meQTL genotypes that are spuriously correlated with age in the discovery EWAS. We also observe that 6.1% of CpGs increase in significance by over one order of magnitude under the conditional analysis, while only 0.023% of CpGs increase by as much in the permutation analysis. Furthermore nine CpG sites (1%) become more significantly associated with age by over two orders of magnitude under the conditional analysis (Figure 8). These results suggest that, for some CpG sites, the genotype state of the true meQTL provides valuable information for characterizing the relationship between methylation level and chronological age.

**Figure 7.**
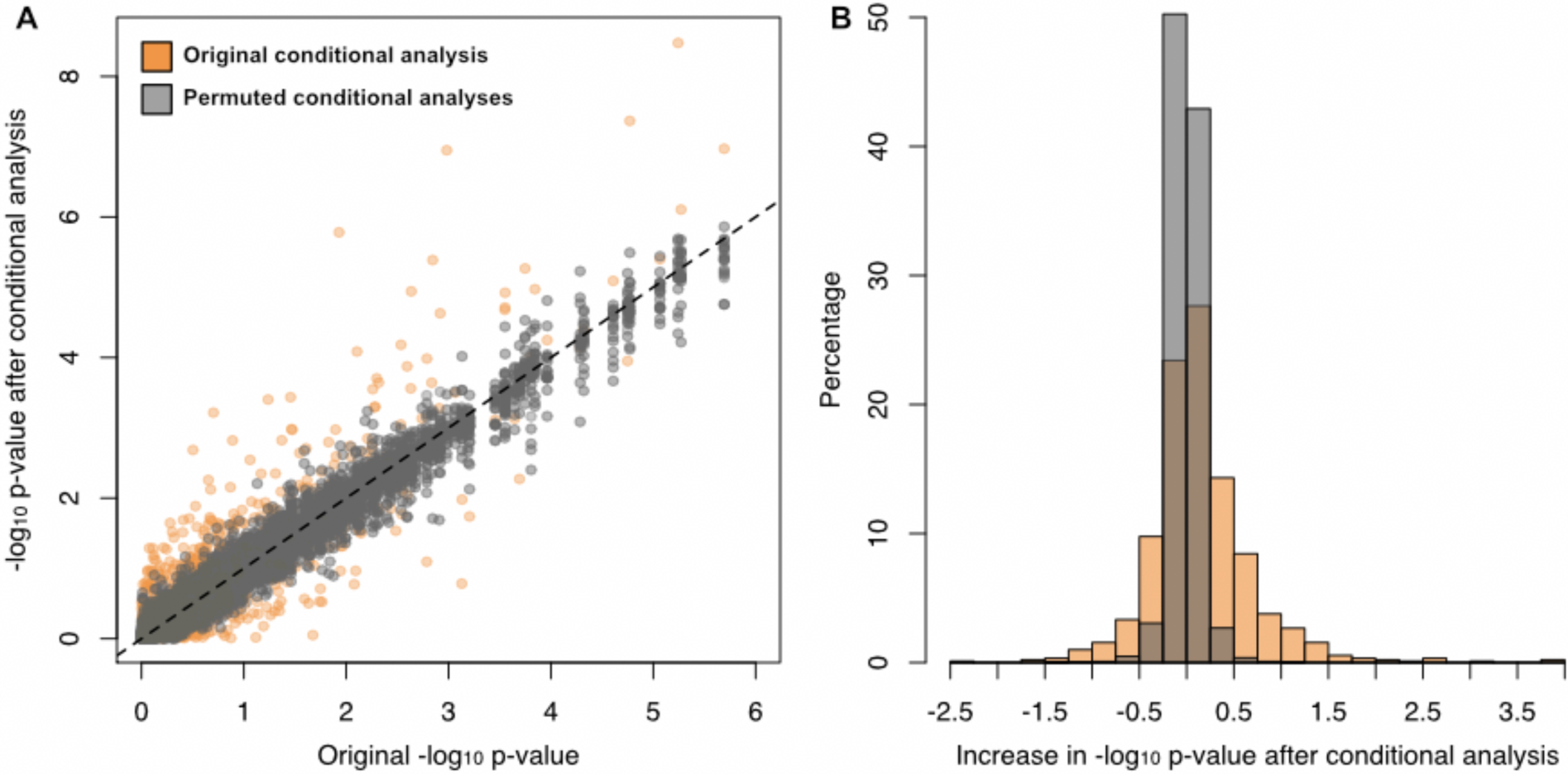
Conditional analysis of meQTL associated a-CpGs. A) The −log_10_ p-values from an epigenome-wide association study on Baka blood are plotted against the −log_10_ p-values from a conditional analysis in which a methylation quantitative trait locus (meQTL) genotype state was included as an additional covariate for 2,842 a-CpGs. B) The distribution of effects of the conditional analysis are depicted as the difference in −log_10_ p-values before and after conditional analysis. The orange points and bars represent the results of the conditional analysis. The grey points and bars represent the results of 100 permutations of the conditional analysis where the CpG-meQTL associations were randomized.

**Figure 8.**
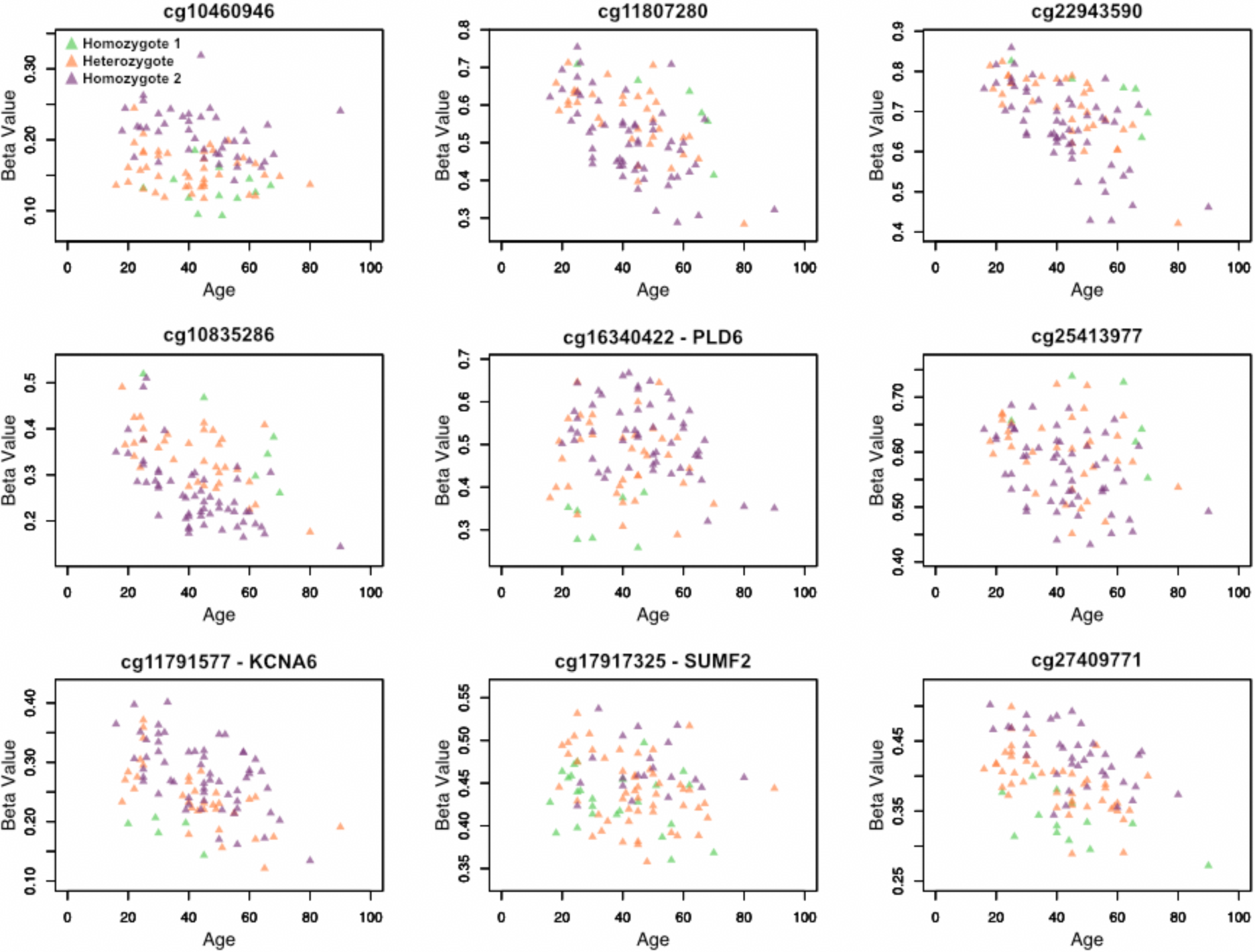
Scatterplots of a-CpGs with associated meQTL genotype states. Scatterplots of beta value and age are shown for the nine CpG sites for which age association improves (i.e. p-value decreases) by over two orders of magnitude when SNP genotype information from a known methylation quantitative trait locus is accounted for in the EWAS. Individuals are coloured by their genotype (homozygous reference, alternative or heterozygous) state, demonstrating genotype-specific trends between methylation and age exist at these CpG sites. Beta values plotted here are not adjusted for the covariates included in each EWAS.

## Discussion

In this study, we investigated patterns of aging in the epigenome across an extended range of human genetic diversity by characterizing the DNA methylation profiles of saliva and whole blood tissues from two modern African hunter-gatherer populations using a large, comprehensive methylation array, which assays over 485,000 CpG sites across the genome. We replicate several of the strongest signals of age-related DNA methylation changes reported in previous studies, including cg16867657 in the gene *ELOVL2,* which supports the utility of this gene as a predictive marker of chronological aging in all humans as was previously suggested by Garagnani et al.^15^. We further demonstrate that this a-CpG replicates strongly in saliva, identifying it independently in both our hunter-gatherer datasets. *ELOVL2* is part of a family of enzymes that are responsible for elongating polyunsaturated fatty acids, whose levels have been shown to decline with chronological age in human skin^66^. It is possible that the continuous life-long increase in methylation of cg16867657 and the *ELOVL2* promoter in general contributes to this trend. It is important to note that this aging biomarker has not been identified in skin tissue itself, but rather in whole blood and white blood cells^2, 19, 25, 43^.

We identified a strong hypomethylation trend with age in the gene D-aspartate oxidase *(DDO),* particularly in saliva tissue. The most significant DDO-annotated CpG site from our study, cg02872426, was found to be significantly age-related in a previous study of an Arab population^25^. The enzyme encoded by DDO deaminates D-aspartic acid, the enantiomer of L-aspartic acid which is the optical form naturally synthesized by biological organisms^67, 68^. Nonenzymatic accumulation of D-aspartic acid is age dependant in living tissues, and is so pronounced in tissues with low turnover that it has been proposed as a biomarker for aging^7^. The role of *DDO* is to eliminate this abnormal version of aspartic acid in proteins and counteract the racemization process and, interestingly, its levels increase in the liver and kidneys with age^67^. The hypomethylation trend we observe in the *DDO* promoter is compatible with these observations and previous age-related methylation studies, and suggests a potential mechanism by which *DDO* expression levels are regulated throughout an organism’s lifetime. Our observations further suggest that the hypomethylation of *DDO,* which may be related to its continued upregulation throughout life, is protective against the effects of age-accumulated protein damage and facilitates ‘healthy’ aging.

A parallel can be drawn between methylation at *DDO* and telomerase reverse transcriptase (*TERT*), their transcriptional regulation, and their function as biomarkers of aging. Shortened telomere length in lymphocytes, a commonly used indicator of biological age, is associated with decreased telomerase levels^6^. Almén et al. observe hypermethylation of *TERT* with age, and speculate that this epigenetic trend is what ultimately underlies the observed trend of telomere shortening with age^18^. Age-related changes in *DDO* methylation may influence gene transcription, but unlike the relationship with *TERT* methylation and telomere shortening, increased levels of *DDO* in older individuals would counteract pathogenic accumulation of abnormal protein. It is tempting to speculate that modulation of *DDO* methylation throughout life is, overall, protective against the detrimental effects of biological aging.

We also identify 107 significant a-CpGs across three EWAS and a meta-analysis that have not, to our knowledge, been reported in any previous study of DNA methylation and aging. 19 of these have high correlation coefficients or strong regression between methylation level and age. Of these, all but cg01519742 are absent from the 27k array and therefore could not be identified in studies using only that technology. However, there exist other difficulties in replicating our results between populations and tissue types, even within our own study. For example, the site cg26559209 exhibits a clear hypomethylation trend in the saliva, but a slight hypermethylation trend in whole blood. This may indicate a tissue-specific pattern of epigenetic aging that is further complicated by the cell-type heterogeneity of whole blood, which, despite bioinformatic correction algorithms, could introduce noise to the aging signal at certain a-CpGs. We also note that many of our novel a-CpGs change only slightly in methylation level over time. The aging signal at these sites may be too weak to be consistently perceptible in other studies, or they may be false positives in our study.

We replicate general trends in the genomic features of a-CpGs, such as the differences in CpG island context of hypermethylated and hypomethylated classes of sites. Previous work on methylation and aging in pediatric cohorts found dramatic changes in methylation patterns occurring during childhood, and that most a-CpGs, both hyper- and hypomethylated, are better modelled by a log-linear relationship between beta value and age^48^. In our study, the methyltyping of Baka children allowed us to observe a similar pattern. However, we also found that hypomethylated a-CpGs were significantly better fit by log-linear models than were hypermethylated a-CpGs. Taken together, this suggests that hyper- and hypomethylated a-CpGs in general are affected differently by aging, and that different biological mechanisms may underlie these epigenetic modifications. It has been generally accepted that the substantial changes in DNA methylation that occur over an organism’s lifetime are mainly a signal of dysregulation of the epigenetic machinery, which ultimately underlies an individual’s age-elevated risk for cellular damage and cancer^4, 69^. In particular, the pervasive hypomethylation with age of CpG sites that lie outside of CpG islands has been spotlighted as an indication of this biological breakdown. Lifestyle and environmental factors can also affect the trajectory of changes in genomic methylation, which can potentially compound or mitigate this risk^9, l8, l9, 70^. However, there are also certain regions of the genome where epigenetic changes appear to be tightly regulated throughout life despite environmental and stochastic variation, and we speculate that these may be protective against the detrimental effects of aging or otherwise adaptive. We hypothesize that *DDO* is an example of a gene that is regulated in such a manner throughout an individual’s life. This is in agreement with the recently proposed conceptual distinction made by Jones et al. between random ‘epigenetic drift’ that may occur due to loss of regulatory control with age and the ‘epigenetic clock’ that is much more precisely correlated with age in humans^5^.

We tested the Horvath model of epigenetic age-prediction built on 353 ‘clock-CpGs’, which were selected only from sites present on both the 27k and 450k arrays, and was trained primarily on European tissue methylation datasets^12^. This model was also tested on chimpanzees in an effort to demonstrate its wide applicability to all human groups, and was found to produce accurate estimates in this closely related species, particularly from whole blood where the correlation between chronological and predicted age was 0.9 and the median error was 1.4 years^12^. However, this type of validation does not account for the possibility of variation in methylation profiles among diverse human populations, potentially resulting from divergent selection on meQTLs or unique environmental or nutritional factors.

We found that the Horvath model does not predict age accurately in our Baka whole blood methylation dataset, and yields an inflated estimate of epigenetic age. We rely on selfreported age in this study, and although it can often be challenging to determine true chronological age in the field, this does not appear to be a driving cause of the inflation observed in Baka blood, as saliva-derived DNA methylation profiles from the same population yield highly accurate estimates of age. As these arrays were run in several batches and separately from our saliva datasets, we speculated that this result might be due to a technical artefact. We explored additional pre-processing pipelines and ComBat batch correction, but could not eliminate the overestimation effect (results not shown). An additional possibility is that the methylation profiles of Baka whole blood are truly epigenetically ‘older’ than European whole blood and the increase in predicted age is biologically meaningful. A recent study of this model found that ancestry-specific differences in epigenetic aging indeed exist and can explain differential mortality rates between ethnic groups^26^. Therefore the population and tissue-specific difference that we observe in the Baka could be driven by genetic background or environmental factors, such as stress, that affect methylation levels at the 353 ‘clock-CpGs’. We performed a sensitivity analysis of the Horvath model to determine if specific CpG sites were disproportionately contributing to the observed overestimation, but did not find any significant results (results not shown). Ultimately, we were unable to determine if systematic over-inflation of predicted ages from our whole blood dataset was due to batch effects, a small variation in the pipelines we used compared to the original study, or a real biological effect.

We found that methylation levels at 901 previously reported a-CpGs are also significantly associated with the genotype state at a *cis* genetic variant. Only eight of these are also significant a-CpGs in our study, and we demonstrate that variation at the associated meQTL is a significant explanatory factor for this lack of replication. By performing a conditional analysis, which accounts for the genotype state of the known meQTL, we were able to recover significant age association in 39 of these 901 CpG sites. For nine CpG sites, including the genotype state at the known meQTL increases the statistical age association by over two orders of magnitude (Figure 8). These sites may prove to be excellent candidates for aging biomarkers or components of an epigenetic age predictor when used in tandem with SNP data, as many of them exhibit large changes in beta value over the measured age range and are strongly correlated with age. The results from our conditional analysis also offer an explanation for the difficulty in replicating a-CpGs from one study to another, namely that differences in the degree of genetic variation at meQTLs confounds the consistent identification of a-CpGs across cohorts, both between and within human populations. Identifying genetic variants that affect a-CpGs is a challenge because the noise introduced by this genetic variability makes it difficult to statistically identify signals of age-related changes in methylation. The approach we use here, which identifies meQTLs at all assayed CpG sites in one cohort and finds overlap with a-CpGs identified in a separate cohort, makes it possible to identify these interactions and recover significant a-CpG association signals.

In this study of African hunter-gatherer DNA methylation patterns, we demonstrate that some CpG methylation changes with age are strongly conserved at specific a-CpGs across genetically diverse human populations and across tissues, and can be confirmed as reliable and universal biomarkers for human aging. We identify 107 novel a-CpGs, which may be useful aging biomarkers. We also observe that genetic variation in a population, particularly at meQTLs, can result in variation in age-related differential DNA methylation. This variation, if uncharacterized or unaccounted for in epigenetic age prediction algorithms, can lead to poor estimates of age in different cohorts and populations. On the other hand, this variation can also be leveraged to improve the precision of age prediction. We conclude that DNA methylation patterns are a promising suite of molecular biomarkers for age across diverse human groups, and that further characterizing these patterns in genetically and ecologically diverse cohorts will facilitate the development of more precise and accurate epigenetic age predictors in the future.

## Acknowledgements

Funding was provided to B.M.H. by a Stanford University CDEHA seed grant (NIH, NIA P30 AG017253-12), to L.Q.-M. by the Institut Pasteur, the CNRS, a CNRS “MIE” (Maladies Infectieuses et Environnement) Grant, and a Foundation Simone & Cino del Duca Research Grant, and to M.S.K. by the Canadian Institute for Advanced Research (CIFAR). M.S.K. is also the Canada Research Chair in Social Epigenetics. We would also like to thank the Working Group of Indigenous Minorities in Southern Africa (WIMSA) and the South African San Institute (SASI) for their encouragement and advice. Finally, we thank the ≠Khomani San and Baka communities in which we have sampled; without their support, this study would not have been possible.

## Accession Numbers

The accession numbers for data used in this paper are in **: [accession number TBD]

---

**Figure S1.**
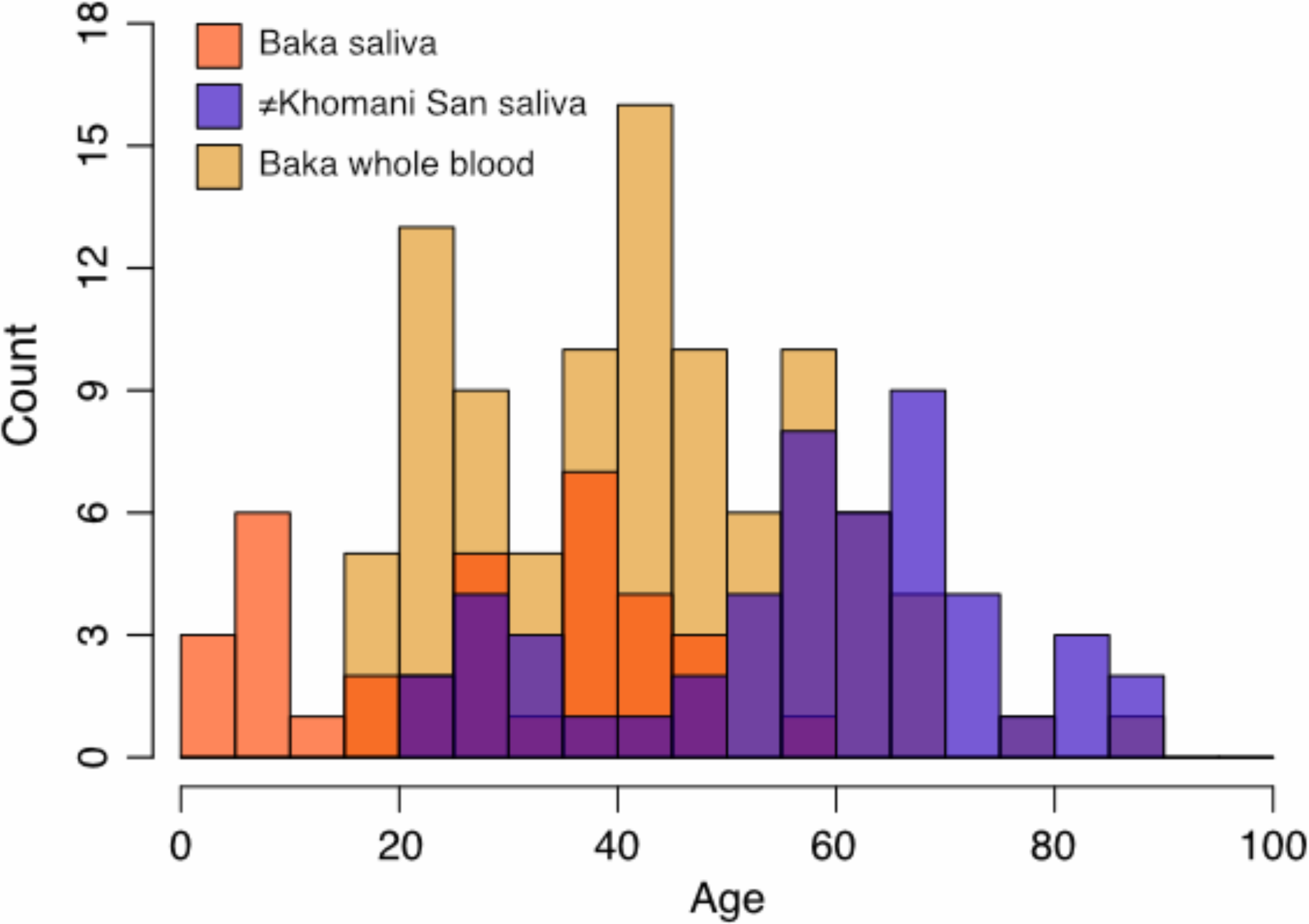
Age structure of the African datasets. Age structure of the Baka saliva, ≠Khomani San saliva and Baka blood datasets based on reported individual ages.

**Figure S2.**
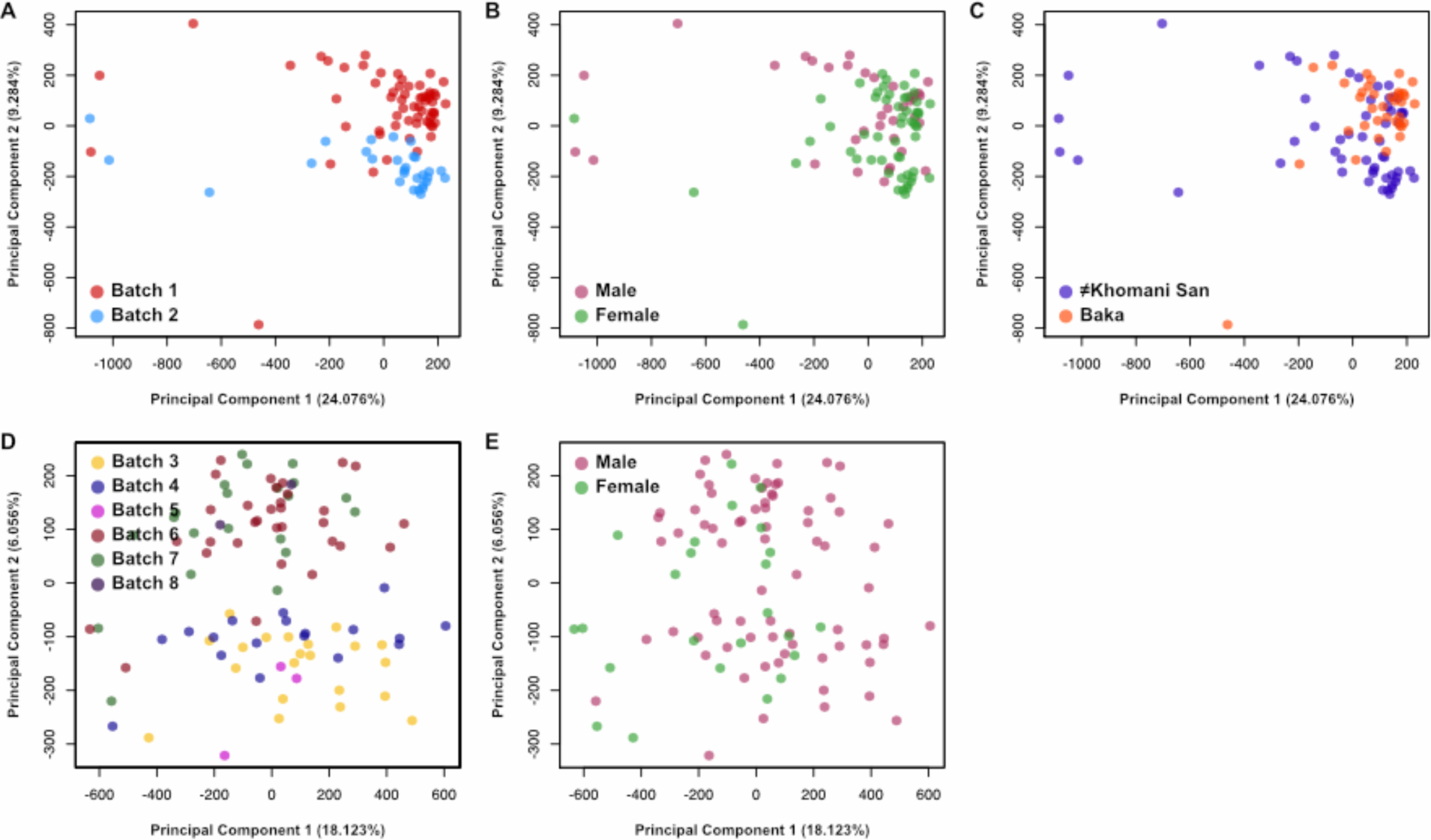
Principal Component Analysis Bi-plots. Principal components analyses (PCA) were performed on both the saliva datasets together and the blood dataset separately. The first two PCs are plotted against each other, coloured by additional variables. The percentage of the variation in the dataset explained by each PC is shown in brackets along the x- and y-axes. Bi-plots of the Baka and ≠Khomani San saliva datasets are coloured by A) batch, B) sex, and C) population. Bi-plots of the Baka blood dataset are coloured by D) batch and E) sex.

**Figure S3.**
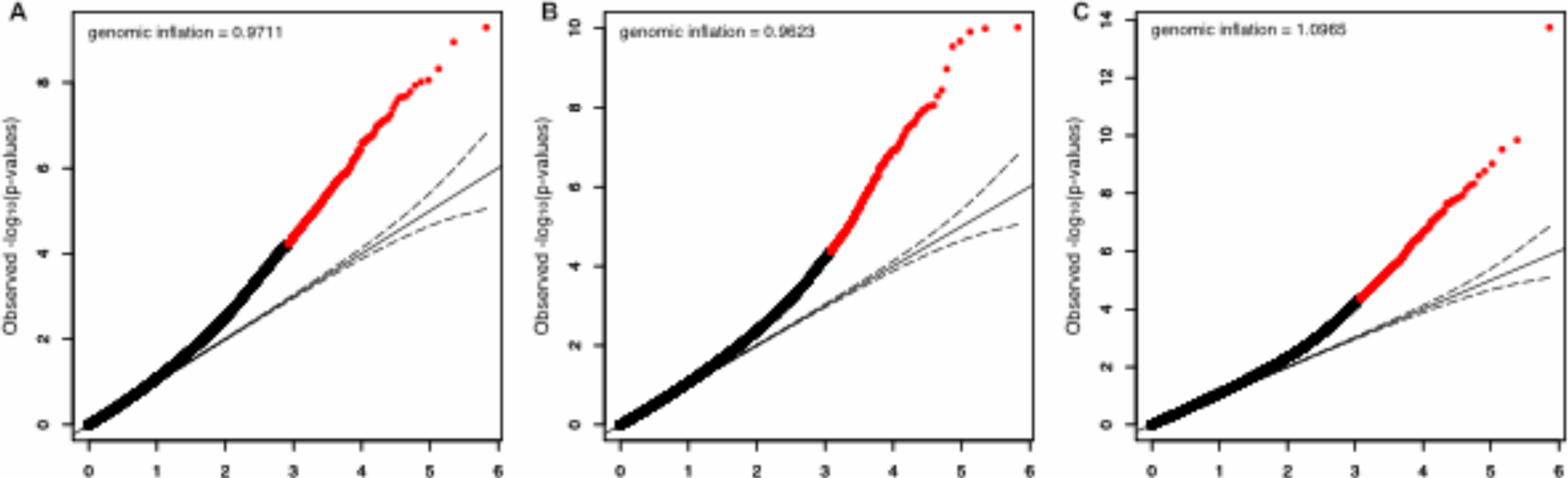
Quantile-quantile plots. The ranked p-values resulting from A) the Baka saliva epigenome-wide association study (EWAS) B) the ≠Khomani San saliva EWAS and C) the Baka blood EWAS are plotted against the expected p-values given no association between methylation level and age. CpG sites that exceeded the Benjamini-Hochberg significance threshold are plotted in red. The dashed curves represent the upper and lower 5% confidence intervals around the null expectation. The genomic inflation factor is shown for each EWAS.

**Figure S4.**
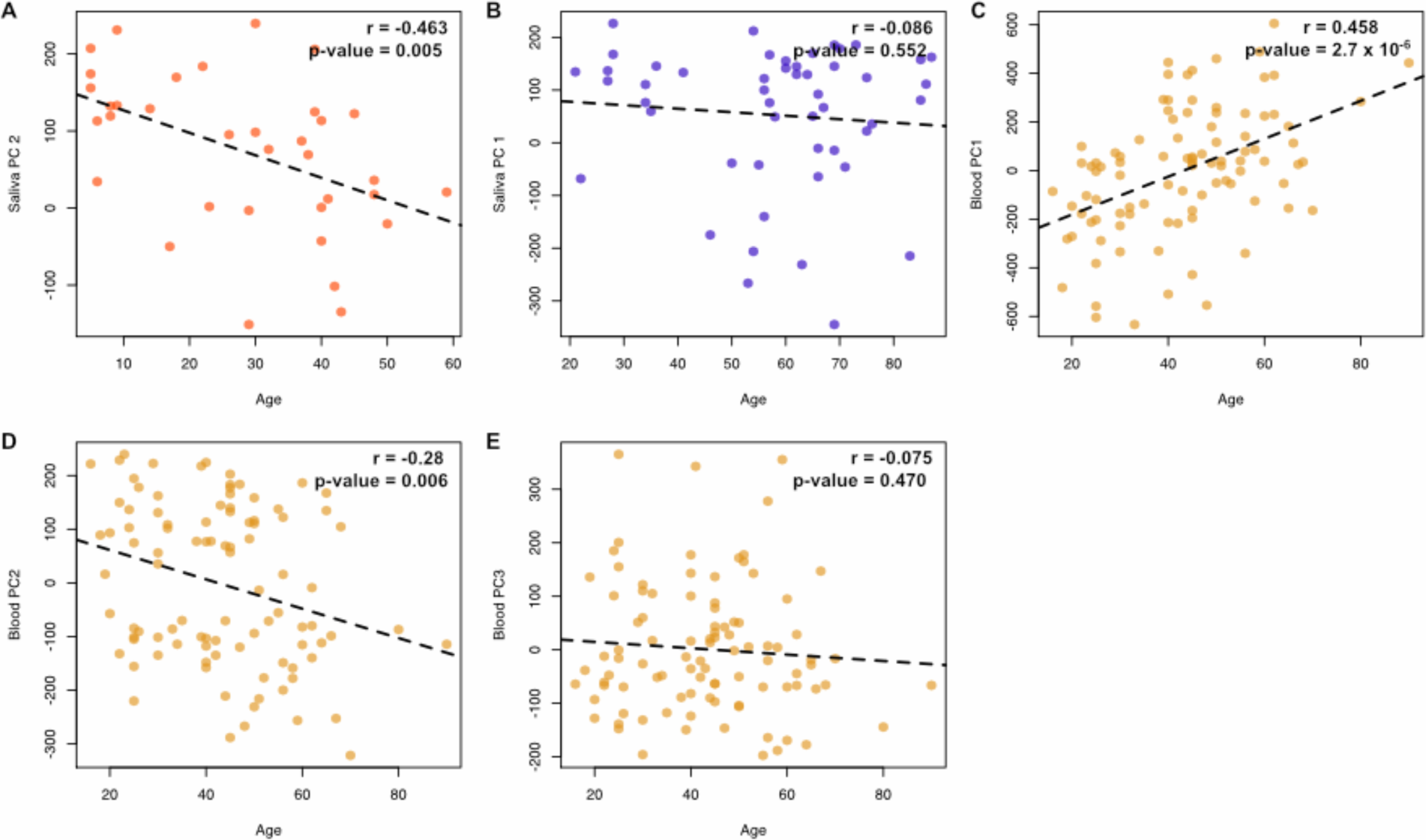
Scatterplot of age and principal component values. Values of the principal components (PC) used as covariates in the epigenome-wide association studies are plotted against chronological age for A) the Baka saliva dataset, B) the ≠Khomani San saliva dataset, and C.E) the Baka whole blood dataset. The dashed lines represent the line of best fit from the linear models of the PC value and age. The Pearson correlation value between age and PC value and p-value of the linear model are shown in each panel.

**Figure S5.**
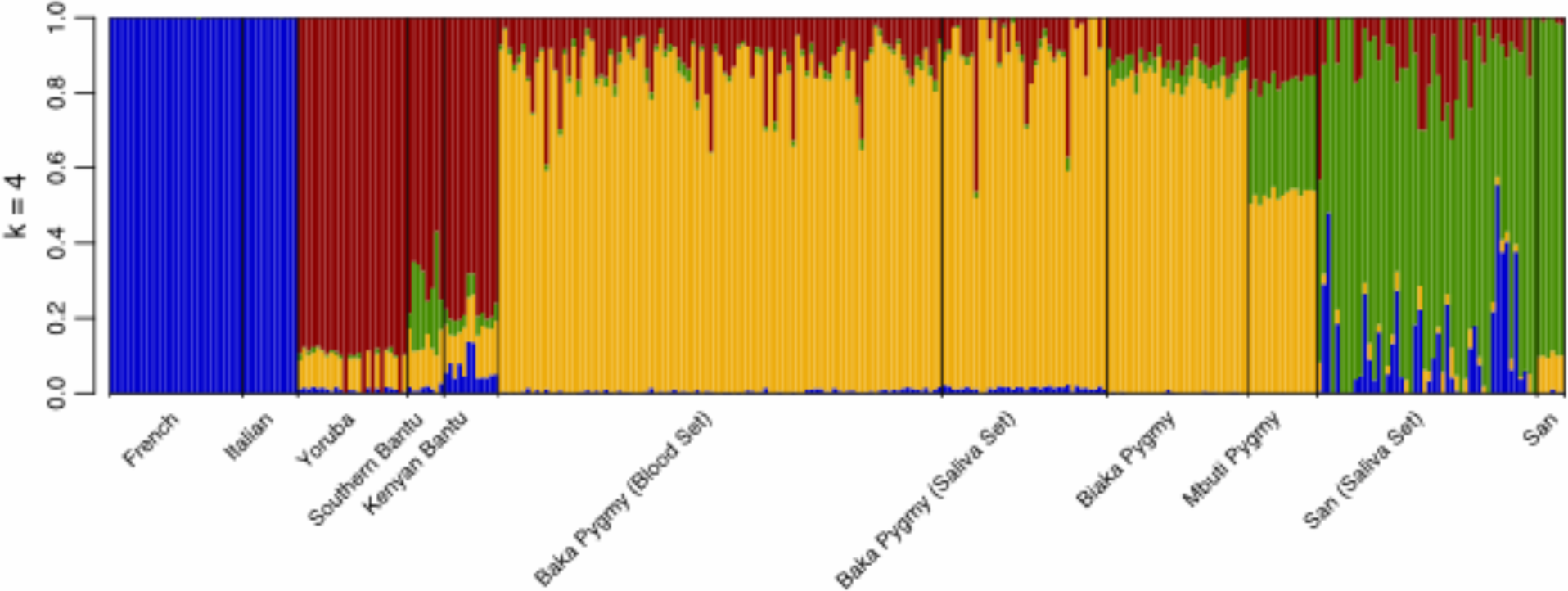
Inferred genome-wide ancestry proportions for Baka and ≠Khomani San samples. Major global ancestry portions for the individuals in the Baka saliva, ≠Khomani San saliva and Baka blood datasets were inferred based on unsupervised clustering of 254,080 SNPs using ADMIXTURE^42^, using additional Illumina 660K SNP array data from a panel of Human Genome Diversity Project (HGDP) populations from Africa and Europe^41^. We focus on k=4 ancestries for the Baka Pygmies and ≠Khomani San, concordant with previous observations^35, 39, 40^.

**Figure S6.**
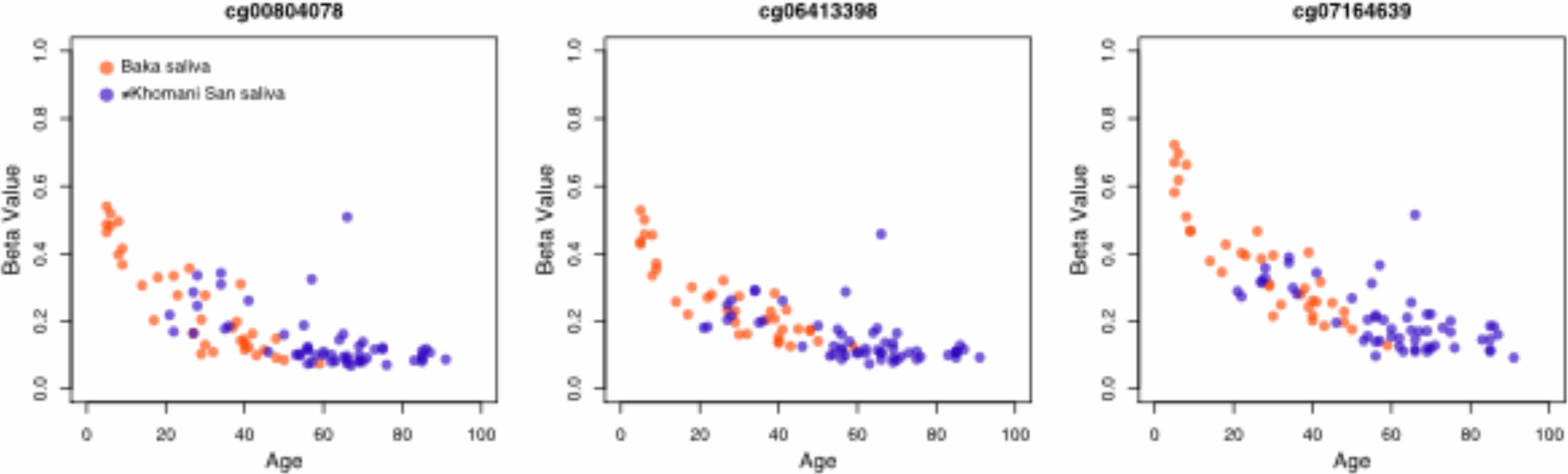
Hypomethylation with age of CpG sites in the gene D-aspartate oxidase. The methylation level is plotted against age for three additional sites in the gene D-aspartate oxidase (*DDO*) that exhibit age-associated hypomethylation at a relaxed significance threshold of p < 0.001 in the two saliva datasets. Beta values plotted here are not adjusted for the covariates included in each EWAS.

## References

1 Hannum, G., Guinney, J., Zhao, L., Zhang, L., Hughes, G., Sadda, S., Klotzle, B., Bibikova, M., Fan, J.B., Gao, Y., et al. (2013). Genome-wide methylation profiles reveal quantitative views of human aging rates. Mol. Cell 49, 359–367.

2 Johansson, A., Enroth, S., and Gyllensten, U. (2013). Continuous aging of the human DNA methylome throughout the human lifespan. PLoS One 8, e67378.

3 Heyn, H., Li, N., Ferreira, H.J., Moran, S., Pisano, D.G., Gomez, a., Diez, J., Sanchez-Mut, J.V., Setien, F., Carmona, F.J., et al. (2012). Distinct DNA methylomes of newborns and centenarians. Proc. Natl. Acad. Sci. 109, 10522–10527.

4 Teschendorff, A.E., West, J., and Beck, S. (2013). Age-associated epigenetic drift: Implications, and a case of epigenetic thrift? Hum. Mol. Genet. 22, 7–15.

5 Jones, M.J., Goodman, S.J., and Kobor, M.S. (2015). DNA methylation and healthy human aging. Aging Cell 14, 924–932.

6 Blasco, M.A. (2007). Telomere length, stem cells and aging. Nat. Chem. Biol. 3, 640–649.

7 Helfman, P.M., and Bada, J.L. (1975). Aspartic acid racemization in tooth enamel from living humans. Proc. Natl. Acad. Sci. 72, 2891–2894.

8 Simm, A., Nass, N., Bartling, B., Hofmann, B., Silber, R.E., and Navarrete Santos, A. (2008). Potential biomarkers of ageing. Biol. Chem. 389, 257–265.

9 Li, Y., Daniel, M., and Tollefsbol, T.O. (2011). Epigenetic regulation of caloric restriction in aging. BMC Med. 9, 98.

10 Holly, A.C., Melzer, D., Pilling, L.C., Henley, W., Hernandez, D.G., Singleton, A.B., Bandinelli, S., Guralnik, J.M., Ferrucci, L., and Harries, L.W. (2013). Towards a gene expression biomarker set for human biological age. Aging Cell 12, 324–326.

11 Meissner, C., and Ritz-Timme, S. (2010). Molecular pathology and age estimation. Forensic Sci. Int. 203, 34–43.

12 Horvath, S. (2013). DNA methylation age of human tissues and cell types. Genome Biol. 14, R115.

13 Bocklandt, S., Lin, W., Sehl, M.E., Sánchez, F.J., Sinsheimer, J.S., Horvath, S., and Vilain, E. (2011). Epigenetic predictor of age. PLoS One 6, e14821.

14 Weidner, C.I., Lin, Q., Koch, C.M., Eisele, L., Beier, F., Ziegler, P., Bauerschlag, D.O., Jockel, K.-H., Erbel, R., Mühleisen, T.W., et al. (2014). Aging of blood can be tracked by DNA methylation changes at just three CpG sites. Genome Biol. 15, R24.

15 Garagnani, P., Bacalini, M.G., Pirazzini, C., Gori, D., Giuliani, C., Mari, D., Di Blasio, A.M., Gentilini, D., Vitale, G., Collino, S., et al. (2012). Methylation of ELOVL2 gene as a new epigenetic marker of age. Aging Cell 11, 1132–1134.

16 Breitling, L.P., Yang, R., Korn, B., Burwinkel, B., and Brenner, H. (2011). Tobacco-smoking-related differential DNA methylation: 27K discovery and replication. Am. J. Hum. Genet. 88, 450–457.

17 Grönniger, E., Weber, B., Heil, O., Peters, N., Stäb, F., Wenck, H., Korn, B., Winnefeld, M., and Lyko, F. (2010). Aging and chronic sun exposure cause distinct epigenetic changes in human skin. PLoS Genet. 6, e1000971.

18 Almén, M.S., Nilsson, E.K., Jacobsson, J.A., Kalnina, I., Klovins, J., Fredriksson, R., and Schiöth, H.B. (2014). Genome-wide analysis reveals DNA methylation markers that vary with both age and obesity. Gene 548, 61–67.

19 Vandiver, A.R., Irizarry, R.A., Hansen, K.D., Garza, L.A., Runarsson, A., Li, X., Chien, A.L., Wang, T.S., Leung, S.G., Kang, S., et al. (2015). Age and sun exposure-related widespread genomic blocks of hypomethylation in nonmalignant skin. Genome Biol. 16, 1–15.

20 Bell, J.T., Pai, A.A., Pickrell, J.K., Gaffney, D.J., Pique-Regi, R., Degner, J.F., Gilad, Y., and Pritchard, J.K. (2011). DNA methylation patterns associate with genetic and gene expression variation in HapMap cell lines. Genome Biol. 12, R10.

21 Gentilini, D., Mari, D., Castaldi, D., Remondini, D., Ogliari, G., Ostan, R., Bucci, L., Sirchia, S.M., Tabano, S., Cavagnini, F., et al. (2013). Role of epigenetics in human aging and longevity: Genome-wide DNA methylation profile in centenarians and centenarians’ offspring. Age 35, 1961–1973.

22 Heyn, H., Moran, S., Hernando-Herraez, I., Sayols, S., Gomez, A., Sandoval, J., Monk, D., Hata, K., Marques-Bonet, T., Wang, L., et al. (2013). DNA methylation contributes to natural human variation. Genome Res. 23, 1363–1372.

23 Fraser, H.B., Lam, L.L., Neumann, S.M., and Kobor, M.S. (2012). Population-specificity of; human DNA methylation. Genome Biol. 13, R8.

24 Fagny, M., Patin, E., Macisaac, J.L., Rotival, M., Flutre, T., Jones, M.J., Siddle, K.J., Quach, H., Harmant, C., Lisa, M., et al. The epigenomic landscape of African rainforest hunter-gatherers and farmers. 1–34.

25 Zaghlool, S.B., Al-Shafai, M., Al Muftah, W.A., Kumar, P., Falchi, M., and Suhre, K. (2015). Association of DNA methylation with age, gender, and smoking in an Arab population. Clin. Epigenetics 7, 1–12.

26 Horvath, S., Gurven, M., Levine, M.E., Trumble, B.C., Kaplan, H., Allayee, H., Ritz, B.R., Chen, B., Lu, A.T., Rickabaugh, T.M., et al. (2016). An epigenetic clock analysis of ' race/ethnicity, sex, and coronary heart disease. Genome Biol. 17, 171.

27 Florath, I., Butterbach, K., Müller, H., Bewerunge-hudler, M., and Brenner, H. (2014). Cross > sectional and longitudinal changes in DNA methylation with age: An epigenome-wide analysis revealing over 60 novel age-associated CpG sites. Hum. Mol. Genet. 23, 1186–1201.

28 Rakyan, V.K., Down, T.A., Thorne, N.P., Flicek, P., Kulesha, E., Gräf, S., Tomazou, E.M., Bäckdahl, L., Johnson, N., Herberth, M., et al. (2008). An integrated resource for genome-wide identification and analysis of human tissue-specific differentially methylated regions (tDMRs). 1518–1529.

29 Byun, H.M., Siegmund, K.D., Pan, F., Weisenberger, D.J., Kanel, G., Laird, P.W., and Yang, A.S. (2009). Epigenetic profiling of somatic tissues from human autopsy specimens identifies ' tissue and individual-specific DNA methylation patterns. Hum. Mol. Genet. 18, 4808–4817.

30 Illingworth, R., Kerr, A., Desousa, D., Helle, J., Ellis, P., Stalker, J., Jackson, D., Clee, C., Plumb, R., Rogers, J., et al. (2008). A novel CpG island set identifies tissue-specific methylation at developmental gene loci. PLoS Biol. 6, e22.

31 Christensen, B.C., Houseman, E.A., Marsit, C.J., Zheng, S., Wrensch, M.R., Wiemels, J.L., Nelson, H.H., Karagas, M.R., Padbury, J.F., Bueno, R., et al. (2009). Aging and environmental exposures alter tissue-specific DNA methylation dependent upon CpG island context. PLoS Genet. 5, e1000602.

32 Verdu, P., and Destro-Bisol, G. (2012). African Pygmies, what’s behind a name? Hum. Biol. 84, 1–10.

33 Veeramah, K.R., Wegmann, D., Woerner, A., Mendez, F.L., Watkins, J.C., Destro-Bisol, G., Soodyall, H., Louie, L., and Hammer, M.F. (2012). An early divergence of KhoeSan ancestors from those of other modern humans is supported by an ABC-based analysis of autosomal resequencing data. Mol. Biol. Evol. 29, 617–630.

34 Verdu, P., Austerlitz, F., Estoup, A., Vitalis, R., Georges, M., Théry, S., Froment, A., Le Bomin, S., Gessain, A., Hombert, J.M., et al. (2009). Origins and genetic diversity of pygmy hunter-gatherers from Western Central Africa. Curr. Biol. 19, 312–318.

35 Henn, B.M., Gignoux, C.R., Jobin, M., Granka, J.M., Macpherson, J.M., Kidd, J.M., Rodriguez-Botigue, L., Ramachandran, S., Hon, L., Brisbin, A., et al. (2011). Hunter-gatherer genomic diversity suggests a southern African origin for modern humans. Proc. Natl. Acad. Sci. 108, 5154–5162.

36 Price, M.E., Cotton, A.M., Lam, L.L., Farré, P., Emberly, E., Brown, C.J., Robinson, W.P., and Kobor, M.S. (2013). Additional annotation enhances potential for biologically-relevant analysis of the Illumina Infinium HumanMethylation450 BeadChip array. Epigenetics Chromatin 6, 4.

37 Dedeurwaerder, S., Defrance, M., Bizet, M., Calonne, E., Bontempi, G., and Fuks, F. (2013). A comprehensive overview of Infinium HumanMethylation450 data processing. Brief. Bioinform.

38 Maksimovic, J., Gordon, L., and Oshlack, A. (2012). SWAN: Subset-quantile within array normalization for Illumina Infinium HumanMethylation450 BeadChips. Genome Biol. 13, R44.

39 Uren, C., Kim, M., Martin, A.R., Bobo, D., Gignoux, C.R., Helden, D. Van, Möller, M., Hoal, E.G., and Henn, B.M. (2016). Fine-scale human population structure in southern Africa reflects ecological boundaries. bioRxiv 038729.

40 Patin, E., Katherine, J.S., Guillaume, L., Hélène, Q., Harmant, C., Becker, N., Froment, A., Régnault, B., Lemée, L., Gravel, S., et al. (2014). The impact of agricultural emergence on the genetic history of African rainforest hunter-gatherers and agriculturalists. Nat. Commun. 5, 3163.

41 Li, J.Z., Absher, D.M., Tang, H., Southwick, A.M., Casto, A.M., Ramachandran, S., Cann, H.M., Barsh, G.S., Feldman, M., Cavalli-Sforza, L.L., et al. (2008). Worldwide human relationships inferred from genome-wide patterns of variation. Science 319, 1100–1104.

42 Alexander, D.H., Novembre, J., and Lange, K. (2009). Fast model-based estimation of ancestry in unrelated individuals. Genome Res. 19, 1655–1664.

43 Steegenga, W.T., Boekschoten, M.V., Lute, C., Hooiveld, G.J., De Groot, P.J., Morris, T.J., Teschendorff, A.E., Butcher, L.M., Beck, S., and Müller, M. (2014). Genome-wide age-related changes in DNA methylation and gene expression in human PBMCs. Age 36, 1523–1540.

44 Houseman, E.A., Accomando, W.P., Koestler, D.C., Christensen, B.C., Marsit, C.J., Nelson, H.H., Wiencke, J.K., and Kelsey, K.T. (2012). DNA methylation arrays as surrogate measures of cell mixture distribution. BMC Bioinformatics 13, 86.

45 Jaffe, A.E., and Irizarry, R.A. (2014). Accounting for cellular heterogeneity is critical in epigenome-wide association studies. Genome Biol. 15, R31.

46 Evangelou, E., and Ioannidis, J.P.A. (2013). Meta-analysis methods for genome-wide association studies and beyond. Nat. Rev. Genet. 14, 379–389.

47 Akaike, H. (1974). A new look at the statistical model identification. IEEE Trans. Autom.

48 Alisch, R.S., Barwick, B.G., Chopra, P., Myrick, L.K., Satten, G.A., Conneely, K.N., and Warren, S.T. (2012). Age-associated DNA methylation in pediatric populations. Genome Res. 22, 623–632.

49 Bell, J.T., Tsai, P.C., Yang, T.P., Pidsley, R., Nisbet, J., Glass, D., Mangino, M., Zhai, G., Zhang, F., Valdes, A., et al. (2012). Epigenome-wide scans identify differentially methylated regions for age and age-related phenotypes in a healthy ageing population. PLoS Genet. 8, e1002629.

50 Cruickshank, M.N., Oshlack, A., Theda, C., Davis, P.G., Martino, D., Sheehan, P., Dai, Y., Saffery, R., Doyle, L.W., and Craig, J.M. (2013). Analysis of epigenetic changes in survivors of preterm birth reveals the effect of gestational age and evidence for a long term legacy. Genome Med. 5, 96.

51 Rakyan, V.K., Down, T.A., Maslau, S., Andrew, T., Yang, T.P., Beyan, H., Whittaker, P., McCann, O.T., Finer, S., Valdes, A.M., et al. (2010). Human aging-associated DNA hypermethylation occurs preferentially at bivalent chromatin domains. Genome Res. 20, 434439.

52 Teschendorff, A.E., Menon, U., Gentry-Maharaj, A., Ramus, S.J., Weisenberger, D.J., Shen, H., Campan, M., Noushmehr, H., Bell, C.G., Maxwell, A.P., et al. (2010). Age-dependent DNA methylation of genes that are suppressed in stem cells is a hallmark of cancer. Genome Res. 20, 440–446.

53 Fernández, A.F., Bayón, G.F., Urdinguio, R.G., Toraño, E.G., Cubillo, I., García-Castro, J., and Delgado-Calle, J. (2015). H3K4me1 marks DNA regions hypomethylated during aging in human stem and differentiated cells. Genome Res. 25, 27–40.

54 Kananen, L., Marttila, S., Nevalainen, T., Jylhävä, J., Mononen, N., Kähönen, M., Raitakari, O.T., Lehtimäki, T., and Hurme, M. (2016). Aging-associated DNA methylation changes in middle-aged individuals: the Young Finns study. BMC Genomics 17, 103.

55 Marttila, S., Kananen, L., Häyrynen, S., Jylhävä, J., Nevalainen, T., Hervonen, A., Jylhä, M., Nykter, M., and Hurme, M. (2015). Ageing-associated changes in the human DNA methylome: genomic locations and effects on gene expression. BMC Genomics 16, 179.

56 Xu, Z., and Taylor, J.A. (2014). Genome-wide age-related DNA methylation changes in blood and other tissues relate to histone modification, expression and cancer. Carcinogenesis 35, 356–364.

57 Wilhelm-Benartzi, C.S., Koestler, D.C., Karagas, M.R., Flanagan, J.M., Christensen, B.C., Kelsey, K.T., Marsit, C.J., Houseman, E.A., and Brown, R. (2013). Review of processing and analysis methods for DNA methylation array data. Br. J. Cancer 109, 1394–1402.

58 Jarvis, J.P., Scheinfeldt, L.B., Soi, S., Lambert, C., Omberg, L., Ferwerda, B., Froment, A., Bodo, J.-M., Beggs, W., Hoffman, G., et al. (2012). Patterns of ancestry, signatures of natural selection, and genetic association with stature in Western African pygmies. PLoS Genet. 8, e1002641.

59 Quintana-Murci, L., Quach, H., Harmant, C., Luca, F., Massonnet, B., Patin, E., Sica, L., Mouguiama-Daouda, P., Comas, D., Tzur, S., et al. (2008). Maternal traces of deep common ancestry and asymmetric gene flow between Pygmy hunter-gatherers and Bantu-speaking farmers. Proc. Natl. Acad. Sci. U. S. A. 105, 1596–1601.

60 Pickrell, J.K., Patterson, N., Barbieri, C., Berthold, F., Gerlach, L., Güldemann, T., Kure, B., Mpoloka, S.W., Nakagawa, H., Naumann, C., et al. (2012). The genetic prehistory of southern Africa. Nat. Commun. 3, 1143.

61 Balding, D.J., and Nichols, R.A. (1995). A method for quantifying differentiation between populations at multi-allelic loci and its implications for investigating identity and paternity. Genetica 96, 3–12.

62 Kang, H.M., Zaitlen, N.A., Wade, C.M., Kirby, A., and Heckerman, D. (2008). Efficient Control of Population Structure in Model Organism Association Mapping. 1723, 1709–1723.

63 Zbieć-Piekarska, R., Spólnicka, M., Kupiec, T., Makowska, ͹., Spas, A., Parys-Proszek, A., Kucharczyk, K., Płoski, R., and Branicki, W. (2015). Examination of DNA methylation status of the ELOVL2 marker may be useful for human age prediction in forensic science. Forensic Sci. Int. 14, 161–167.

64 Ali, O., Cerjak, D., Kent, J.W., James, R., Blangero, J., Carless, M.A., and Zhang, Y. (2015). An epigenetic map of age-associated autosomal loci in northern European families at high risk for the metabolic syndrome. Clin. Epigenetics 7, 12.

65 Smith, A.K., Kilaru, V., Kocak, M., Almli, L.M., Mercer, K.B., Ressler, K.J., Tylavsky, F.A., and Conneely, K.N. (2014). Methylation quantitative trait loci (meQTLs) are consistently detected across ancestry, developmental stage, and tissue type. BMC Genomics 15, 145.

66 Kim, E.J., Kim, M., Jin, X., Oh, J., Kim, J.E., and Chung, J.H. (2010). Skin aging and photoaging alter fatty acids composition, including 11, 14, 17-eicosatrienoic acid, in the epidermis of human skin. J. Korean Med. Sci. 25, 980–983.

67 D’Aniello, A., D’Onofrio, G., Pischetola, M., D’Aniello, G., Vetere, A., Petrucelli, L., and Fisher, G.H. (1993). Biological role of D-amino acid oxidase and D-aspartate oxidase: Effects of D-amino acids. J. Biol. Chem. 268, 26941–26949.

68 Ritz-Timme, S., and Collins, M.J. (2002). Racemization of aspartic acid in human proteins. Ageing Res. Rev. 1, 43–59.

69 Jaenisch, R., and Bird, A. (2003). Epigenetic regulation of gene expression: how the genome integrates intrinsic and environmental signals. Nat. Genet. 33 Suppl, 245–254.

70 Zannas, A.S., Arloth, J., Carrillo-Roa, T., Iurato, S., Röh, S., Ressler, K.J., Nemeroff, C.B., Smith, A.K., Bradley, B., Heim, C., et al. (2015). Lifetime stress accelerates epigenetic aging in an urban, African American cohort: relevance of glucocorticoid signaling. Genome Biol. 16, 266.

